# The effects of reliable social feedback on language learning: insights from EEG and pupillometry

**DOI:** 10.1101/2025.02.04.636399

**Authors:** Ana Zappa, Polina Osokina, Xim Cerda-Company, David Cucurell, Maria Mateu, Antoni Rodriguez-Fornells

**Author notes:** Corresponding author: Ana Zappa, Department of Cognition, Development and Educational Psychology, University of Barcelona, Barcelona, Spain.

## Abstract

Language learning is often a social process, and social feedback may play a motivational role. We examined the neurophysiological correlates of word learning with feedback varying in reliability and social content. Participants associated novel auditory words with objects and received social (video clips) or symbolic (static images) feedback. In a forced-choice task, participants learned to associate novel auditory words with known objects and received feedback that was either Social Reliable (correct), Social Unreliable (random), or Symbolic Reliable (correct). Post-training behavioral performance was better for words learned with social and symbolic reliable feedback. Stimulus-preceding negativity (SPN) and late positive complex (LPC) ERP amplitudes, as well as pupil dilation, showed differences as a function of feedback reliability and social content. In the reliable conditions, before feedback, SPN amplitude grew as learning progressed, likely due to the expectation of receiving positive feedback. During feedback, LPC amplitude for positive feedback diminished as learning progressed but not for negative feedback, which was likely consistently used for context updating. These effects were not observed for unreliable feedback, probably because its value was not used for updating information. Pupillometry results corroborated these findings, showing greater dilation for negative vs positive feedback in reliable conditions. Finally, when feedback was social, processing was associated with more frontal activation and behavioral performance was closely correlated with both ERP and pupillometry results. Overall, our findings show differential processing of feedback depending on its informational and social content, advancing our understanding of how social and cognitive processes interact to shape word learning.

## Introduction

Language learners frequently receive social feedback, informing them as to their success or failure in language performance (e.g., pronouncing a newly learned word), which they use to update linguistic knowledge and adapt future behavior (Mackey & Sachs, 2012). Evaluative social feedback, expressed through verbal or non-verbal reactions (e.g., praise, encouragement, disapproval), plays a key role in social learning (Sobczak & Bunzenk, 2023; Zonca et al., 2021; for a review, see Ho et al., 2017). Beyond its informational value, social feedback has motivational and emotional significance, serving as a source of reward or disapproval (Triconi & DePasque, 2017), which reinforces adaptive behavior. Despite the importance of social feedback in gating learning, its impact on language learning remains poorly understood. The current study investigated the neurocognitive mechanisms underlying the processing of social feedback during word learning in adults, as well as how different types of feedback affect word acquisition.

Learning to associate novel labels to existing semantic concepts, or word learning, is a fundamental aspect of language learning. Feedback, particularly when reliable, facilitates this process by reinforcing the association between an auditory word the corresponding object (or image). The effectiveness of feedback can be explained by associative theories of predictive learning, which emphasize the role of prediction errors in driving attention and updating associations. When feedback outcomes deviate from learners’ expectations, prediction errors occur, prompting updates to mental representations and a gradual reduction in error signals over time (Luque et al., 2012). Importantly, cues that generate prediction errors tend to attract more attention than those confirming correct predictions (Pearce and Hall, 1980; Wills et al., 2007).

A key factor in feedback processing is reliability, which influences how learners monitor performance and evaluate feedback. Reliable feedback has been shown to enhance both feedback anticipation and processing compared to unreliable feedback (Ernst & Steinhauser, 2017; Schiffer et al., 2017; Severo et al., 2020; Walentowska et al., 2016). Indeed, learners prioritize information from reliable sources, minimizing the costs associated with acquiring inaccurate or irrelevant feedback (Begus et al., 2016; Koenig & Harris, 2005; Mangardich & Sabbagh, 2018; Sabbagh & Baldwin, 2001; Scofield & Behrend, 2008). This preference for reliable sources is particularly evident during early language acquisition but likely extends to adult word learning. From an evolutionary perspective, selective social learning offers adaptive advantages, ensuring efficient acquisition of relevant knowledge (Henrich & McElreath, 2007; Sperber et al., 2010).

A common form of feedback in language learning is social feedback, which is generally prioritized over non-social information (Wagner et al., 2016; Chevallier et al., 2012; Williams et al., 2019). Its inherent emotional content provides an additional processing advantage, as emotional information is often processed more deeply and rapidly (Hariri et al., 2002; Norris et al., 2004). Positive or negative social feedback may serve as a powerful motivator for word learning by activating reward-motivational systems and enhancing both attention and memory processes (Mani & Ackermann, 2018; Martins et al., 2021). A major difficulty in observing the effect of social feedback on learning is providing participants with ecologically valid stimuli that mimic real-life human feedback. Most previous studies have used either static pictures of faces (Spreckelmeyer et al., 2009; for a review, see Risko et al., 2012) or still images of thumbs up/down (Pfabigan et al., 2017), which lack the inherent temporal dynamics of social feedback. In contrast, videos that show real humans in movement are thought to better reflect real-life interactions, leading to higher emotional arousal and motivational engagement (Khols et al., 2013; Sato & Yoshikawa, 2007).

Feedback-related processes during learning are often examined using event-related potentials (ERPs). The stimulus-preceding negativity (SPN) is thought to reflect feedback expectation and anticipation, capturing motivational, affective, and preparatory processes tied to expected information (Morís et al., 2013; Böcker et al., 2002; Kotani et al., 2003; Masaki et al., 2006; Poli et al., 2007; Glazer et al., 2018; Leon-Cabrera et al., 2024). The late positive complex (LPC or P3b) is associated with cognitive mechanisms such as working memory updating and attention allocation (Donchin & Coles, 1998; Polich, 2007). Enhanced LPC amplitudes have been linked to encoding processes that predict improved subsequent recall and retrieval performance (Batterink & Neville, 2011; Fabiani et al., 1986; Karis et al., 1984; Lemhöfer et la., 2025; Turk et al., 2018).

Another effective measure of feedback processing is pupil dilation (Mathôt, 2018). This psychophysiological response has been shown to index the anticipation of rewards (Schneider et al., 2018), interest (Hess & Polt, 1960), emotional arousal (Bradley et al., 2008) and curiosity for learning new information (Kang et al., 2009; Brod et al., 2019, Theobald et al., 2022). In line with this evidence, pupil dilation can be used as a marker of expectation and anticipation of upcoming feedback. Furthermore, pupil dilation has been widely examined in connection to mental effort (Laeng et al., 2012) and is thought to predict successful learning (Kafkas, 2021, for a review, see Kafkas, 2024), memory load (Kahneman and Beatty, 1966; Vilotijevic & Mathôt, 2023), strength of memory (Otero et al., 2011; Pajkossy & Racsmány, 2019) and recognition (for a metanalysis, see Lapteva & Martarelli, 2024).

In the current study, we tracked the neurocognitive processes underlying word learning with feedback that differed in its reliability and social content. Using a within-subjects design, we examined feedback expectation/anticipation and processing, as well as its impact on novel word acquisition. Positive and negative social feedback was presented via a series of short video clips and symbolic feedback was presented as static images. Participants learned to associate novel auditory words with known objects through a forced-choice task, receiving feedback in three conditions: Social Reliable (correct), Social Unreliable (random), and Symbolic Reliable (correct). A block design ensured participants were aware of the feedback type prior to its onset. Behavioral learning performance was evaluated during training and in a post-training assessment.

Given the social nature of language learning, we hypothesized that reliable social feedback is a relevant and efficient “currency” for motivating word learning. We predicted that prior to feedback, reliable social feedback would result in greater SPN amplitude and pupil dilation, reflecting heightened feedback expectation and anticipation compared to symbolic and unreliable social feedback. During feedback, we predicted that reliable social feedback would evoke greater LPC amplitude and pupil dilation, indicative of enhanced feedback processing and better encoding, particularly for negative feedback. If supported, these findings would suggest that social feedback not only increases feedback expectation but also facilitates deeper processing and encoding, highlighting its role as a critical motivator in second language acquisition.

## 2. Materials and methods

### 2.1. Participants

Thirty-two (22 females, mean age: 23 ± 5 yrs) right-handed Spanish-Catalan bilinguals with no previous knowledge of Hebrew or related languages participated in the study. Sample size was based on previous ERP studies on feedback processing. Participants had normal or corrected-to-normal vision and reported no history of neurological deficits. Three participants were excluded from the EEG analyses due to excessive noise in the EEG signal and two due to technical sound difficulties during the experiment. Out of the remaining 27 participants, 5 were excluded from the pupillometry analyses due to recording issues. Before the experiment, participants gave written consent and were fully informed of the experimental procedure. They received 30€ for their participation in the 2-hour session, 45-minute preparation and 15-minute online post-test. The study was approved by the local ethics committee.

### 2.2. Stimuli

90 grey scale images were selected from the MultiPic standardized set of drawings (Duñabeitina et al., 2018) and 90 matching auditory nouns in Hebrew were recorded by a native speaker. Nouns belonged to six different categories: animals and insects, body parts, food, clothing, household items and tools. Verbal labels were not transparent with the equivalent label in Spanish, Catalan, French, Italian, German or English. The lexicon was separated into three separate lists controlled for word frequency and word length, which contained the same number of words from each category. Different lists were assigned to different feedback conditions for each participant, counterbalanced across participants. Feedback conditions during learning included: Symbolic Reliable feedback, Social Reliable feedback and Social Unreliable feedback. For Symbolic Reliable feedback, participants saw an image of a check (positive) or cross (negative) (Fig.1). Six different videos for each Agent, with a duration of 2 secs, were recorded by two professional actresses (one was Agent 1 and the other Agent 2) who were physically distinctive from each other. In order to provide varied social feedback, social feedback consisted of: thumbs up/down, smile/frown, nod/head-shaking. All visual materials were controlled for isoluminance due to the use of pupillometry. Half of the participants saw Agent 1 in the Social Reliable condition and Agent 2 in the Social Unreliable condition; the other half had the opposite combination. In the Social Reliable and Symbolic Reliable conditions, participants always received 100% correct feedback during training (after selecting the image they thought corresponded to each auditory noun). In the Social Unreliable condition, feedback was 50% correct and 50% incorrect, making it unreliable.

**Figure 1.**
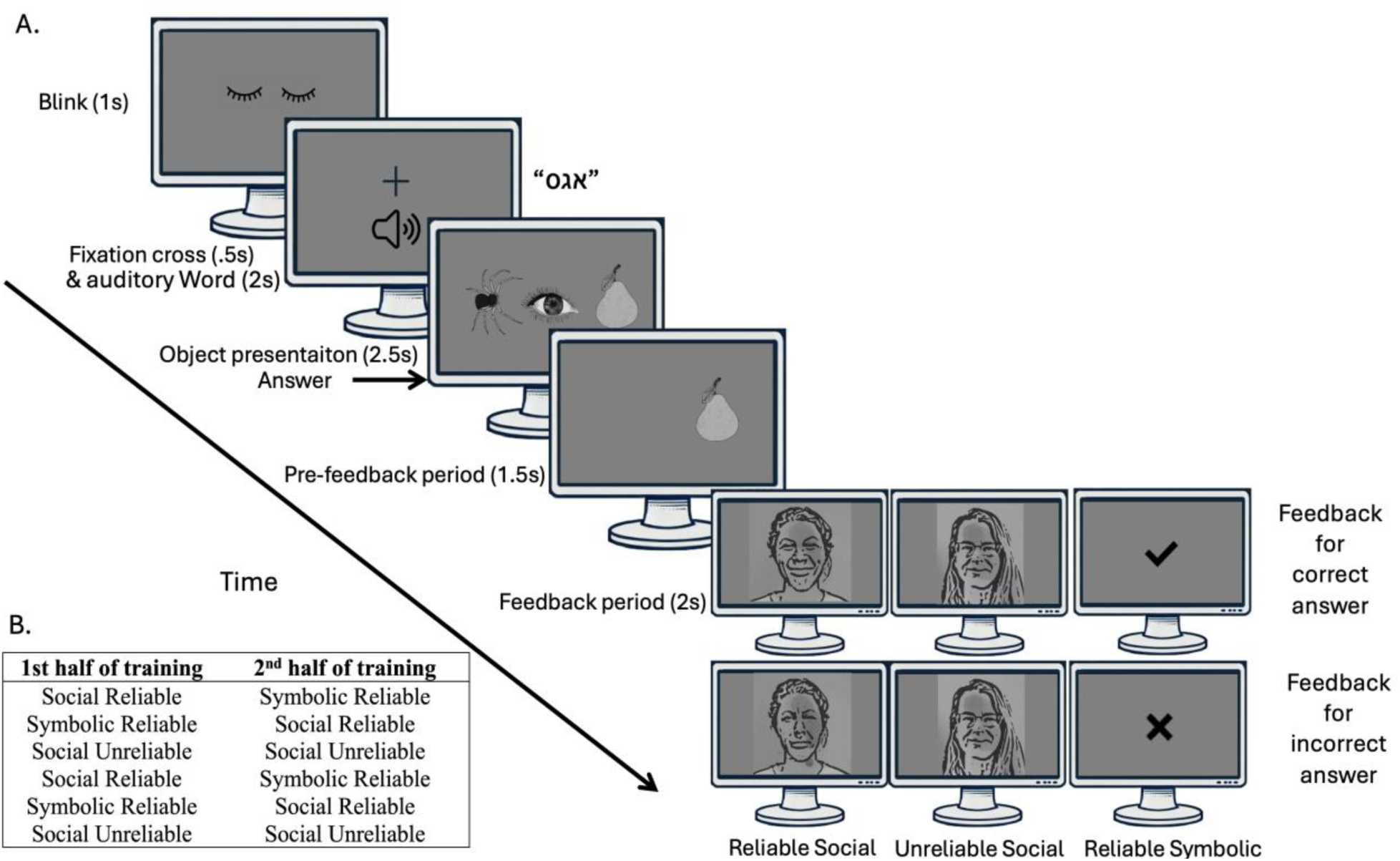
Trial design and block order example. **A**. Trial design: participants heard a novel word, saw three images, selected one of the three using a keyboard and received feedback regarding the accuracy of their selection. In each of the six blocks (2 per condition), they received feedback in one of three conditions: Reliable Social, Unreliable Social or Symbolic. **B.** An example of block order for the first and second half of training.

A separate experiment with 20 participants that were not involved in the current study was conducted to ensure that all feedback was equally perceived as positive or negative and to measure possible processing response timing differences between conditions. Participants were asked rate how positive or negative the feedback was using a 5-point Likert scale. At least 90% of the participants rated 1 (fully negative) for negative feedback, and 5 (fully positive) for positive feedback in all symbolic and social feedbacks. Social feedback was perceived as negative or positive 0.960 (sd=0.130) sec after Symbolic feedback [Symbolic mean=0.624 sec, (sd=0.072 sec), Social Agent 1, mean=1.592 sec, (sd=0.126 sec), Social Agent 2, mean=1.575 sec, (sd=0.134 sec)]. No significant differences were found in response time between negative [mean=1.612 sec, (sd=0.112 sec)] and positive [mean=1.555 sec, (sd=0.140 sec)] social feedback videos, nor negative [mean=0.644 sec, (sd=0.091 sec)] and positive [mean=0.633 sec, (sd=0.062 sec)] symbolic videos.

### 2.3. Procedure and tasks

#### 2.3.1 Training

Participants were briefed on the procedure, capped with an elastic cap containing 32 electrodes and fitted with Pupil Labs glasses (Kassner et al., 2014). They were comfortably seated at a desk situated 60 cm away from a computer screen in a Faraday booth. Participants underwent a pre-training with three auditory word and image pairs not included in the main training. During the pre-training, they were introduced to the two social agents and told that one of them had less teaching experience (Social Unreliable) than the other (Social Reliable) agent. Following this, they underwent a 2-hour training session, including short breaks after each block, during which they were invited to stretch and rest if necessary. After the first half of training (roughly 55 minutes), participants were given a 10-minute break during which the screen was turned off and they could talk to the examinator and have a snack.

The 90 items were separated into three conditions (30 per condition) and 8 blocks. Half of the items (15 in each condition) were learned during the first half of training and the other half during the second half of training (after the 10-minute break). Conditions were separated by blocks and each item was presented 3 times per block, 6 times total. Participants always began with a block of Symbolic Reliable or Social Reliable condition to ensure that they “trusted” the feedback in the beginning of training. The order of the first three blocks was then repeated to avoid different amounts of time between first 3 and second 3 item exposures in the different conditions. An illustration of a trial can be seen in Fig.1a and an example of block order in Fig.1b. A trial began with a blink prompt (1 sec), followed by the presentation of a centered fixation cross which preceded the auditory verb by 500 ms and then remained on the screen during the subsequent presentation of an auditory word in Hebrew over speakers (2 secs). Participants then saw three items: the target, a distractor from the same list and a distractor from another list (2.5 secs). After they selected an image using the 1, 2 and 3 key on the number pad of a keyboard, to indicate the item on the left side (1), middle (2) or right side (3) of the screen, the selected image remained on the screen for 1.5 secs. If participants took longer than 2.5 seconds to answer, the objects disappeared and a message saying “You must answer more quickly” appeared on the screen. Target position was randomly assigned. Following the post-answer 1.5 secs period, participants saw either Social Reliable, Social Unreliable or Symbolic Reliable feedback (2 secs), depending on the block.

#### 2.3.2 Online post-test

All participants performed an online post-test ten days after training. They heard a noun, saw three objects, and were asked to select the object they thought corresponded to the auditory word. Words learned in all three conditions were mixed and presented three times throughout the list, in pseudorandom order. The four items presented consisted of the target, a distractor from the same list, and two distractors from the two other lists.

### 2.4 Data acquisition

#### 2.4.1 Behavioral accuracy and response time

Accuracy and response time was recorded during training and during the online post-test.

#### 2.4.2 EEG

Electroencephalographic (EEG) activity was recorded continuously from the scalp using a BrainAmp DC amplifier (BrainVision Recorder software, Brain Products©). Electrodes were located at 29 standard positions (Fp1/2, Fz, F7/8, F3/4, FC1/2, FC5/6, FCz, Cz, C3/4, T7/8, CP1/2, CP5/6, Pz, P3/4, P7/P8, PO3/4, and Oz). Biosignals were referenced online to the right outer canthus eye electrode and re-referenced offline to the mean of the activity of two electrodes placed on the mastoids. Vertical eye movements were monitored with an electrode at the infraorbital ridge of the right eye. Electrode impedances were kept below 5 kΩ.

#### 2.4.3 Pupillometry

To measure changes in pupil diameter, the participants wore an eye-tracking glasses (Pupil Labs, Kassner et al., 2014). This system monitored the pupil center, providing the absolute pupil diameter and a confidence index, which ranged from 0 (pupil not detected) to 1 (pupil detected with very high certainty; Faraji et al., 2023). The eye-tracker had a non-constant sampling rate of 200 Hz, and it captured the pupil diameter of both eyes separately, using the dark pupil method in reference to a three-dimensional geometric model of the eyeball (Yeung et al., 2021).

### 2.5 Data analyses

#### 2.5.1. Behavioral data analysis

We performed a generalized linear mixed effects model (glmer) to examine accuracy during the last exposure of training and in the post-test. The fixed effects factor was Condition (Social Reliable, Social Unreliable and Symbolic Reliable) and random factors were Participant and Item.

#### 2.5.2. ERP analyses Preprocessing

A noncausal Butterworth high-pass filter was applied to the continuous EEG (half-amplitude cutoff = 0.01 Hz, slope = 12 dB/octave). The electrophysiological signals were digitized at a rate of 500 Hz. Data was processed and filtered using ERPLAB (Lopez-Calderon & Luck 2014). Artifact rejection was performed offline and tailored to each participant. Independent component analysis (ICA) was carried out on the continuous data, for each participant, to exclude artifacts due to blinking. Before ICA, sections of the EEG signal that were highly contaminated with noise were removed. Following ICA rejection, the data was segmented and divided into the separate conditions. A time-window 1500 ms time-locked to the response (and 100 ms before the answer, as a baseline) was used for the SPN analyses. A time-window 2000 ms time-locked feedback onset (and 100 ms before feedback onset, as a baseline) was used for the LPC analyses. Epoch rejection criteria were individually determined using a simple voltage threshold +/− 100μV. None of the participants’ rejection rate exceeded 20%. ERP figures were created using ERPLAB v2021.0.

##### Analysis

The ERP data was modeled using linear mixed effect models for the mean voltage amplitudes in the pre-feedback and feedback time-windows. For the pre-feedback period, the analyses were time-locked to the answer and conducted on the 200-1000 time-period (after the answer), on the following electrodes: CP1, CP2, P3, P4, Pz, PO3, PO4 (Brunia et al., 1988; Brunia et al., 2011). Windows for feedback-related analyses were selected using a comparison between negative and positive feedback, in each condition, using cluster-based permutation tests that performed 50,000 permutations (Maris & Oostenveld, 2007). over the full post-stimulus time window (0 to 1998 ms), across all exposures, on the following electrodes: C3, C4, CP1, CP2, Cz, Oz, P3, P4, Pz, PO3, PO4 (Sun et al., 2024; Yang et al., 2019).

#### 2.5.3. Pupillometry analyses

All the pupillometry preprocessing was performed using Matlab and considering guidelines from Kret & Sjak-Shie, 2019. The preprocessing proceeded as follows: first, data with confidence values below 0.6 were removed (Yeung et al., 2021). Next, the data was filtered to identify and exclude three common types of invalid pupil size samples: dilation speed outliers and edge artifacts, trend-line deviation outliers, and temporally isolated samples. Valid samples were then interpolated using at a high sampling rate (120 Hz) and processed to calculate the mean measure from both eyes. The resulting signal was smoothed using a zero-phase low-pass filter, with a cutoff frequency of 4 Hz (Kret & Sjak-Shie, 2019; Jackson & Sirois, 2009). Finally, external triggers indicating stimuli onsets were used to segment the pupillometry data. Similarly to ERP analyses, for the SPN analysis, epochs of 1.5 sec were time-locked to the answer onset, while for feedback analyses, epochs of 2 sec were time-locked to the feedback onset. All epochs included a −500 ms pre-stimulus time-window used for baseline correction.

Pupil dilation within the analyzed time windows was examined by analyzing the grand-averaged change in the pupil diameter. For both pre- and feedback periods, a linear mixed-effects model was fitted, considering the mean change in pupil diameter as the dependent variable, Condition and Exposure as fixed effects and Participant as a random factor. In the pre-feedback period, we analyzed the time window from 250 ms onwards. To determine the analyzed time window for the post-feedback we preformed a nonparametric cluster-based Monte Carlo simulation analysis with 10,000 permutations (Maris & Oostenveld, 2007). The cluster permutation test compared, with a significance level of p < .05, the change in pupil diameter between positive and negative feedback. For subsequent mixed models analyses, mean change in pupil diameter was averaged across all significant time-windows for the statistical analysis. Finally, the differences in mean pupil diameter changes between positive and negative feedback were correlated with behavioral performance by means of a Pearson correlation.

## 3. Results

### 3.1 Behavioral accuracy and response time

#### 3.1.2 During *training*

We first plotted participants performance during the whole training period. As expected, visual inspection of the learning curves showed faster and ultimately more successful learning for the Social Reliable and Symbolic Reliable conditions compared to the Social Unreliable condition (Fig.2). Visual inspection of the post-test plots also revealed improved performance for the Social Reliable and the Symbolic Reliable conditions compared to the Social Unreliable condition.

**Figure 2.**
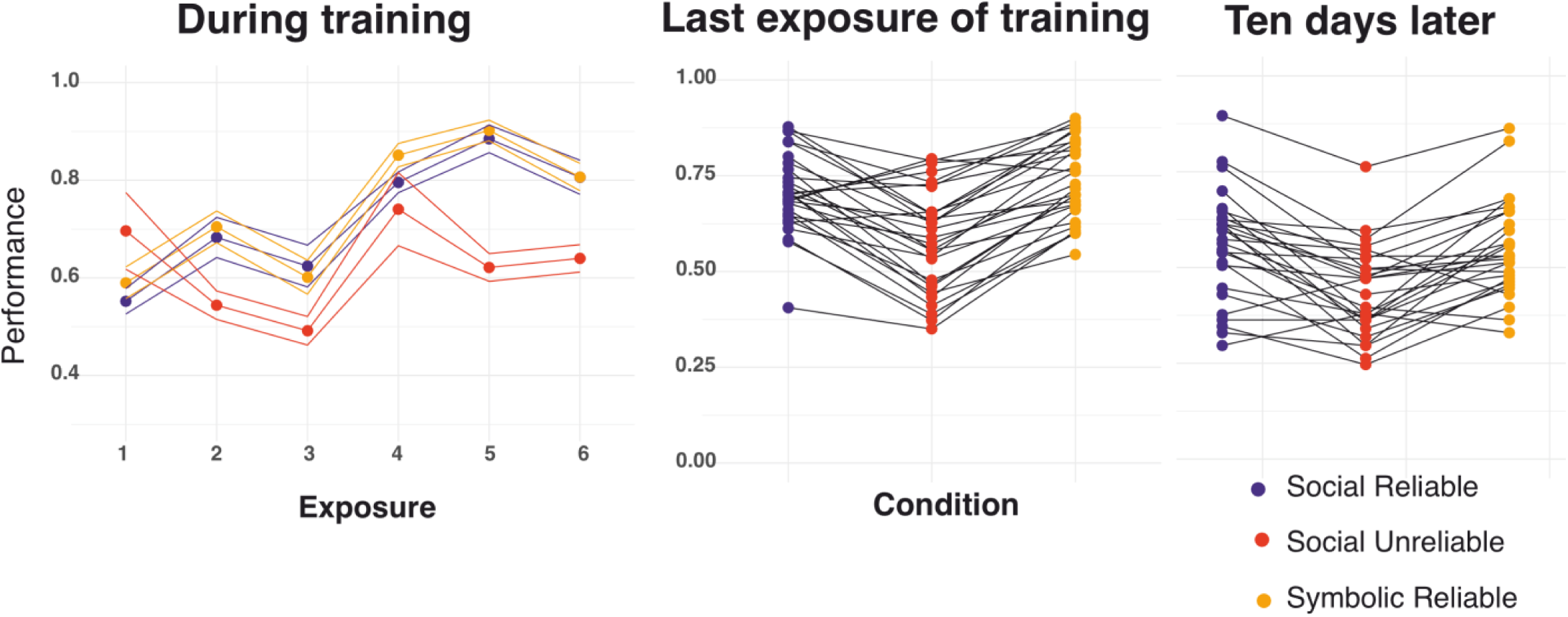
Average accuracy during training over 6 exposures of items with a 95% confidence interval (left), average accuracy per participant at the last exposure of training (middle) and 1-week post-training (right).

##### Behavioral performance during last Block of Learning: Mixed Model results

To measure differences in performance as a function of condition during the last phase of learning (last block of exposure for each item) we performed a general linear mixed effects model that included the fixed factor Condition (Social Reliable, Social Unreliable and Symbolic Reliable), and Participant and Item as random factors. The model [Performance ∼ Condition + (1 | participant) + (1 | item), AIC=2141687.2, LL=-1070840] revealed poorer performance for the Social Unreliable compared to Symbolic reliable (β = −1.22537, se = 0.22865, z = −5.359, p < .001) condition and no differences between the Social Reliable and Symbolic Reliable conditions (β = 0.04336, se = 0.27354, z = −0.159, p =0.874) (Table 1, Fig.2).

**Table 1.** Generalized linear mixed model for behavioral performance during the last block during training.

| Immediate post-test |  |  |  |  |
| --- | --- | --- | --- | --- |
|  | Estimate | SE | z value | p |
| Intercept (Symbolic) | 1.93415 | 0.24565 | 7.874 | 3.45e-15 *** |
| Reliable | 0.04336 | 0.27354 | 0.159 | 0.874 |
| Unreliable | -1.22537 | 0.22865 | -5.359 | 8.37e-08 *** |
| Random effects |  |  |  |  |
|  | Variance | SD |  |  |
| Item (Intercept) | 0.3772 | 0.6141 |  |  |
| Participant (Intercept) | 0.9207 | 0.9596 |  |  |
| Model: Performance ~ Condition + (1 \| participant) + (1 \| target) |  |  |  |  |
| Sampling Units | Number of total observations = 2520 |  |  |  |
|  | Number of participants = 29; Number of items = 90 |  |  |  |
| AIC | BIC | LL |  |  |
| 2141687 | 2141730 | -1070840 |  |  |

#### 3.1.2 Behavioral performance ten days post-training

To measure differences in retention of auditory word-image association as a function of condition in which items were learned during training, 1 week post training, we performed a general linear mixed effects model that included the fixed factor Condition, and Participant and Item as random factors. Once again, the model [Performance ∼ Condition + (1|participant) + (1|item), AIC=6185.2, LL=-3087.6] revealed poorer performance for the Social Unreliable compared to Symbolic Reliable (β = −0.58887, se = 0.08510, z = −6.920, p < .001) and Social Reliable (β = −0.03100, se = 0.08556, z = −0.362, p =0.717102). No differences emerged between the Symbolic Reliable and Social Reliable conditions (β = 0.03100 se = 0.08556, z = 0.362, p =0.717) (Table 2, Fig.1).

**Table 2.** Generalized linear mixed model for behavioral performance 1-week post-training.

| 1-week post-training test |  |  |  |  |
| --- | --- | --- | --- | --- |
|  | Estimate | SE | z value | p |
| Intercept (Symbolic) | 0.46484 | 0.13311 | 3.492 | 0.000479 *** |
| Reliable | -0.03100 | 0.08556 | -0.362 | 0.717102 |
| Unreliable | -0.58887 | 0.08510 - | -6.920 | 4.51e-12 *** |
| Random effects |  |  |  |  |
|  | Variance | SD |  |  |
| Item (Intercept) | 0.3924 | 0 .6264 |  |  |
| Participant (Intercept) | 0.2677 | 0 .5174 |  |  |
| Model: Performance ~ Condition + (1 \| participant) + (1 \| item) |  |  |  |  |
| Sampling Units | Number of total observations = 4809 |  |  |  |
|  | Number of participants = 27; Number of items = 90 |  |  |  |
| AIC | BIC | LL |  |  |
| 6185.2 | 6217.6 | -3087.6 |  |  |

### 3.2 EEG

#### 3.2.1 Pre-feedback

##### 3.2.1.1 Visual inspection

We compared the ERP waveforms preceding feedback, per exposure, in the three conditions. Visual inspection of ERPs in the pre-feedback window revealed a progressive development of a sustained negativity (SPN) with exposure, in the Symbolic Reliable and Social Reliable conditions. However, no evidence of an exposure effect was observed in the Social Unreliable condition. The scalp distributions in Fig.3. represents the difference waveform between the 6^th^ and 1^st^ exposure, during the 200-1000 time-period (after the answer). In the Social Reliable and Symbolic Reliable conditions, the topography is centro-posterior (Fig.3).

**Figure 3.**
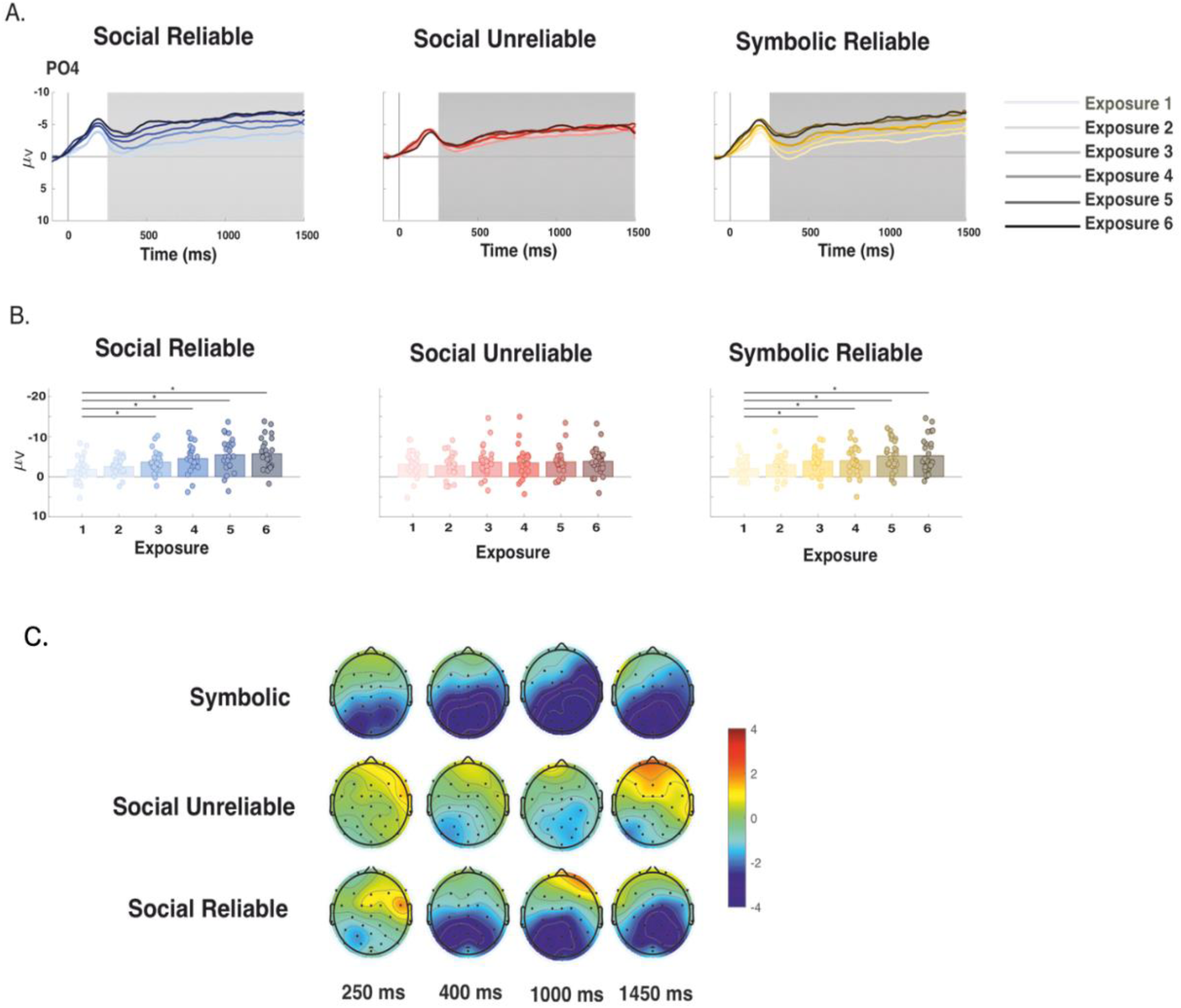
Pre-feedback ERP analysis by exposure **A.** ERPs for different learning conditions at electrode PO4, time-locked to the participant’s answer. For each plot, the lightest line of the given color scheme represents the 1^st^ exposure and gets progressively darker for each exposure. The grey area shows the analyzed time window (200-1000 msec). **B.** Barplots represent the grand average amplitude in the analyzed time window while dots represent each participant’s mean amplitude, for the given exposure (1-6). Asterisks represent significant differences compared to the 1^st^ exposure (p<.05). **C.** Topographies represent the difference waveforms between 6^th^ and 1^st^ exposure. The difference waveforms considered the grand average ERPs between 250ms and 1500 before answer with 150 ms intervals).

##### 3.2.1.2 Linear mixed-effects models

We ran linear mixed effects models, which included Condition and Exposure, and compared the mean voltage amplitudes in the 200-1500 ms post answer onset, at a subset of centro-parietal electrodes (CP1, CP2, P3, P4, Pz, PO3, PO4). The forward selection strategy was used to select the model that best fitted the pre-feedback data. Model 0 included only the random intercept. We then added the predictor Condition (Symbolic Reliable, Social Reliable and Social Unreliable), then the predictor Exposure (Exposure 1,2,3,4,5 and 6) and finally their interaction (Table 3). The final lmer model [MV ∼ condition*exposure + (1|participant) + (1|electrode), p= < 2.2e-16 ***, AIC= 12822, LL=-6390.1] revealed significant interactions, including that of Exposure by Condition (Table 4). Subsequent post-hoc comparisons, corrected for multiple comparisons using the False Discovery Rate (FDR), further delineated these effects. Pair-wise comparisons showed that, in the Social Reliable condition, the SPN amplitude was greater for exposures 2, 3, 4, 5 and 6 compared to exposure 1 (Table 5). The same was true for the Symbolic Reliable condition (Table 5). This interaction was absent in the Unreliable Social condition (Table 5).

**Table 3.** Model comparison for the evaluation pre-feedback EEG.

**Pre-Feedback**
| **Sampling Units** |  | Number of total observations = 2916<br>Number of participants = 27<br>Number of electrodes = 6 |  |  |  |  |  |  |  |  |  |
| --- | --- | --- | --- | --- | --- | --- | --- | --- | --- | --- | --- |
| **Model specification** | **Model name** | **Nested/**<br>**simpler**<br>**model** | **Fixed**<br>**effects**<br>**added** | **Random effects** |  | **Model fit** |  |  |  | **LRT test**<br>**against nested** |  |
|  |  |  |  | **Participants** | **Electrodes** | **AIC** | **BIC** | **LL** | **df** | **df** | **p** |
| RE only | Null | --- | --- | Intercepts | Intercepts | 13486 | 13510 | -<br>6739.1 | 4 | --- | --- |
| FE main<br>effect | Condition | Null | Condition | “ | “ | 13466 | 13502 | -<br>6727.3 | 6 | 2 | 7.405e-06<br>*** |
| FE main<br>effects | Exposure | Condition | Exposure | “ | “ | 12822 | 12948 | -<br>6390.1 | 21 | 15 | < 2.2e-16<br>*** |

RE= Random effects; FE= Fixed effects

**Table 4.** Linear fixed effects models for pre-feedback EEG.

| **Fixed effects** |  |  |  |  |  |
| --- | --- | --- | --- | --- | --- |
|  | **Estimate** | **SE** | **df** | **t** | **p** |
| (Intercept) | -1.2682 | 0.6607 | 35.8748 | -1.919 | 0.06292 |
| ConditionSymbolic | -0.4117 | 0.2338 | 2883.9929 | -1.761 | 0.07839 |
| ConditionUnreliable | -1.6919 | 0.2338 | 2883.9929 | -7.235 | 5.92E-13*** |
| Exposure2 | -1.1741 | 0.2338 | 2883.9929 | -5.021 | 5.45E-07*** |
| Exposure3 | -2.2553 | 0.2338 | 2883.9929 | -9.645 | < 2e-16 *** |
| Exposure4 | -3.1483 | 0.2338 | 2883.9929 | -13.464 | < 2e-16 *** |
| Exposure5 | -3.8405 | 0.2338 | 2883.9929 | -16.424 | < 2e-16 *** |
| Exposure6 | -3.8491 | 0.2338 | 2883.9929 | -16.461 | < 2e-16 *** |
| ConditionSymbolic:Exposure2 | -0.1523 | 0.3307 | 2883.9929 | -0.46 | 0.64522 |
| ConditionUnreliable:Exposure2 | 1.6694 | 0.3307 | 2883.9929 | 5.048 | 4.74E-07*** |
| ConditionSymbolic:Exposure3 | 0.1607 | 0.3307 | 2883.9929 | 0.486 | 0.62708 |
| ConditionUnreliable:Exposure3 | 1.756 | 0.3307 | 2883.9929 | 5.31 | 1.18E-07*** |
| ConditionSymbolic:Exposure4 | 1.0637 | 0.3307 | 2883.9929 | 3.217 | 1.31E-03*** |
| ConditionUnreliable:Exposure4 | 2.8111 | 0.3307 | 2883.9929 | 8.501 | < 2e-16 *** |
| ConditionSymbolic:Exposure5 | 0.51 | 0.3307 | 2883.9929 | 1.542 | 0.12309 |
| ConditionUnreliable:Exposure5 | 3.3136 | 0.3307 | 2883.9929 | 10.02 | < 2e-16 *** |
| ConditionSymbolic:Exposure6 | 0.6013 | 0.3307 | 2883.9929 | 1.818 | 0.0691 |
| ConditionUnreliable:Exposure6 | 3.1619 | 0.3307 | 2883.9929 | 9.562 | < 2e-16 *** |

| **Random effects** |  |  |
| --- | --- | --- |
|  | **Variance** | **SD** |
| Participant (Intercept) | 9.3526 | 3.0582 |
| Electrode (Intercept) | 0.3769 | 0.6139 |

| **Model fit** |  |  |
| --- | --- | --- |
| **R<sup>2</sup>** | **Marginal** | **Conditional** |
|  | 0.0781102 | 0.7116275 |

Key: p-values for fixed effects calculated using Satterthwaites approximations. Confidence intervals have been calculated using the Wald method. Model equation: $MV \sim \text{condition} * \text{correct before} + (1 | \text{participant}) + (1 | \text{electrode})$

**Table 5.** Post-hoc comparisons for pre-feedback EEG.

**Social Reliable**
| **contrast** | **estimate** | **SE** | **df** | **z.ratio** | **p value** |
| --- | --- | --- | --- | --- | --- |
| Exposure1-Exposure2 | 1,1741 | 0,235 | 2901 | 5,006 | <.0001* |
| Exposure2-Exposure3 | 1,08115 | 0,235 | 2901 | 4,61 | <.0001* |
| Exposure3-Exposure4 | 0,89306 | 0,235 | 2901 | 3,808 | 0,0002* |
| Exposure4-Exposure5 | 0,69214 | 0,235 | 2901 | 2,951 | 0,0034* |
| Exposure5-Exposure6 | 0,00867 | 0,235 | 2901 | 0,037 | 0,9705 |

**Social Unreliable**
| **contrast** | **estimate** | **SE** | **df** | **z.ratio** | **p value** |
| --- | --- | --- | --- | --- | --- |
| Exposure1-Exposure2 | -0,49526 | 0,235 | 2901 | -2,112 | 0,0652 |
| Exposure2-Exposure3 | 0,99453 | 0,235 | 2901 | 4,241 | 0,0001* |
| Exposure3-Exposure4 | -0,16209 | 0,235 | 2901 | -0,691 | 0,5294 |
| Exposure4-Exposure5 | 0,18965 | 0,235 | 2901 | 0,809 | 0,5287 |
| Exposure5-Exposure6 | 0,16037 | 0,235 | 2901 | 0,684 | 0,5294 |

**Symbolic**
| **contrast** | **estimate** | **SE** | **df** | **z.ratio** | **p value** |
| --- | --- | --- | --- | --- | --- |
| Exposure1-Exposure2 | 1,32637 | 0,235 | 2901 | 5,656 | <.0001* |
| Exposure2-Exposure3 | 0,76821 | 0,235 | 2901 | 3,276 | 0,0013* |
| Exposure3-Exposure4 | -0,00995 | 0,235 | 2901 | -0,042 | 0,9662 |
| Exposure4-Exposure5 | 1,24579 | 0,235 | 2901 | 5,312 | <.0001* |
| Exposure5-Exposure6 | -0,08263 | 0,235 | 2901 | -0,352 | 0,7764 |

We then compared exposures 1 to 2, 2 to 3, 3 to 4, 4 to 5 and 5 to 6, to verify whether the SPN increased incrementally across the learning session (Table 5). In the Social Reliable condition, the increase in SPN was significant between exposures 1 and 2 (β =1.1741. se = 0.235, z = 5.006, p<.0001), 2 and 3 (β =1.08115, se = 0.235, z = 4.61, p<.0001), 3 and 4 (β =0.89306, se = 0.235, z = 3.808, p=0,0002), and 4 and 5 (β =0.69214, se = 0.235, z = 2.951, p=0.0034). In the Social Unreliable condition, the increase in SPN was only significant between exposures 2 and 3 (β =0.99453, se = 0.235, z = 4.241, p=0.0001). In the Symbolic Reliable condition, the increase in SPN was significant between exposures 1 and 2 (β =1.32637, se = 0.235, z = 5.656, p<.0001), 2 and 3 (β =10.76821, se = 0.235, z = 3.276, p=0.0013), and 4 and 5 (β =1.24579, se = 0.235, z = 3.276, p<.0001). These results show a clear pattern of increasing SPN in the Social Reliable condition, with 4 out of 5 changes showing increasing negativity as the learning session progressed. This pattern was less clear in the Symbolic Reliable condition, with 3 out of 5 of the changes showing a significant increasing negativity but the other two showing a (non-significant) diminution. Finally, this pattern was not at all visible in the Social Unreliable condition, where only one significant increase was seen between exposures 2 and 3 (β =0.99453, se = 0.235, z = 4.241, p<.0001) (Fig.3).

#### 3.2.2 During feedback

##### 3.2.2.1 Visual inspection

We compared the ERP waveforms after feedback presentation, separately for positive and negative feedback in each condition. Visual inspection revealed a temporal difference in feedback processing such that the Symbolic Reliable condition showed greater positivity in an earlier time window (around 200-700 ms) whereas as the Social Reliable and Social Unreliable conditions showed greater positivity in a later time window (around 1200-2000 ms), likely due to the differential nature of the feedback (Fig.4). While the symbolic feedback was immediately presented after participants answered, in the Social conditions, the videos took longer to fully reveal the valence of the feedback (see Section 2.2). We also saw greater positivity in the feedback window for negative compared to positive feedback for all conditions, but the difference appeared greater for the Symbolic Reliable and Social Reliable conditions compared to the Social Unreliable condition. Finally, visual inspection of the topo plots of the difference wave (negative minus positive feedback in the feedback window) revealed less positivity for the Social Unreliable condition (in line with ERP results). It also showed a wider, more frontal, distribution for the Social Reliable compared to the Symbolic Reliable condition (Fig.4).

**Figure 4.**
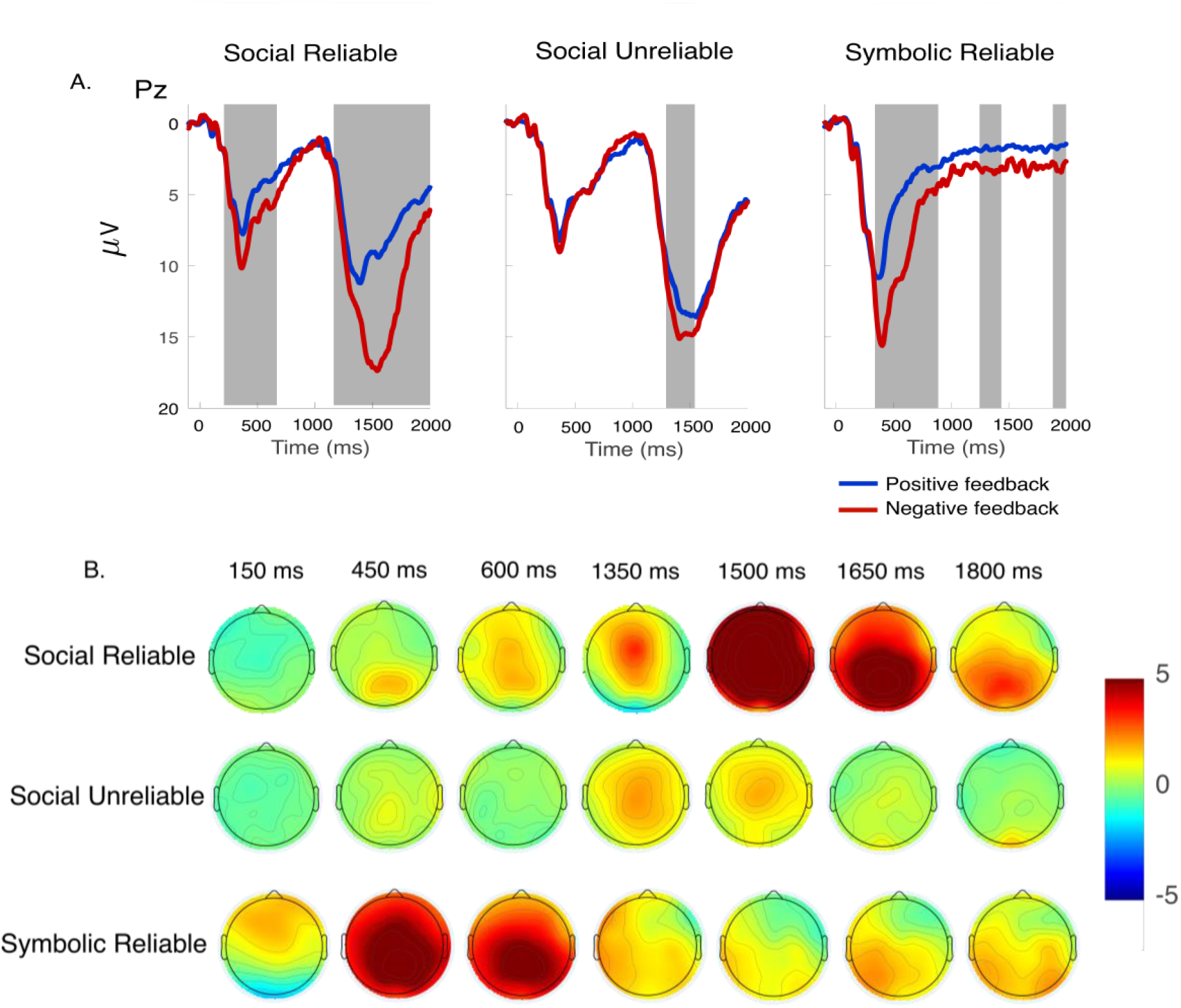
Positive vs negative feedback. **A.** Feedback ERPs at electrode Pz for different learning conditions time-locked to feedback onset. The blue line corresponds to positive feedback and the red line to negative feedback. The grey areas show the time periods during which cluster-based permutation tests revealed significant differences between amplitudes associated with Positive and Negative feedback (p<.05). **B.** Topographies represent the difference waveforms between negative and positive feedback from 250 to 2000 ms with 50 ms intervals.

##### 3.2.2.2 Cluster-based permutation tests

Following visual inspection, we used cluster-based permutation tests to observe possible differences between negative and positive feedback and to determine the time windows to be used for subsequent mixed models analyses (which included the factor Exposure). Significant clusters (shown in grey, Fig.4) emerged in the Social Reliable condition between 314 - 746 ms (tmass: 902.22, p<0.001) and 1210 −1998 ms (tmass: 2263.90, p<0.001) post-stimulus. For the Unreliable Social condition differences emerged between 1290 and 1540 ms (tmass: 390.48. p=0.0044) post-stimulus. Considering the behavioral timing differences between Social and Symbolic stimuli processing (see Section 2.2.), we used the second window for the late positivity analysis in the Social Reliable condition (which overlapped with that of the Social Unreliable condition). For the Symbolic Reliable condition, differences emerged between 338-888 ms (tmass: 1647.90, p<0.001), 1246-1436 ms (tmass: 227.02, p=.0289), and 1882-1998 ms (tmass:155.90, p=.0471) post-stimulus. Given that the effect was going in the same direction for all three windows we collapsed them for mixed model statistical analysis (Fig.4).

##### 3.2.2.3 Topographical analysis

Given the visual differences between the Social Reliable and Symbolic Reliable difference wave (negative minus positive feedback) topo plots in this feedback window, we ran a linear mixed effects models to explore topographical differences. We first ran the model [MV ∼ condition*electrode + (1|participant)] in the mean voltage amplitudes in the 100 ms time window surrounding the peak amplitude in both conditions. Several interactions with Electrode emerged and we hence separated the electrodes into three separate ROIs: frontal (F3, F4, F7, F8, Fp1, Fp2, Fz, FC1, FC2, FC5, FC6, FCZ), central (C3, C4, CP1, CP2, CP5, CP6, Cz) and parietal (Pz, P3, P4, P7, P8, PO3, PO4, Oz) region. We ran a second model that included ROI as a fixed factor [MV ∼ condition*ROI + (1|participant)]. The model [MV ∼ condition*ROI + (1|participant)] revealed significant interactions of Condition by ROI (Table 6).

**Table 6.**
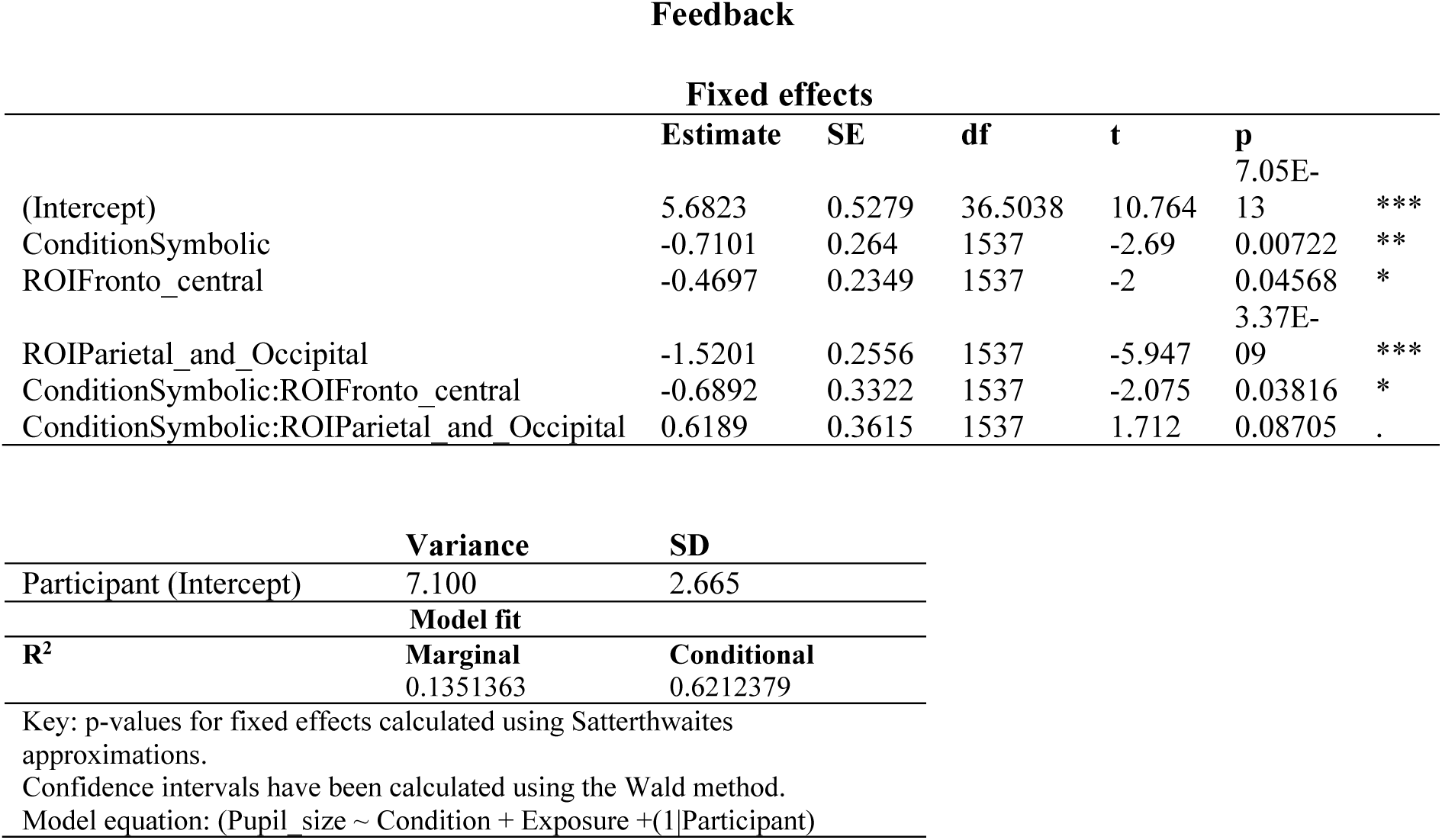
Linear fixed effects model for topographical differences of Feedback differences waves.

Subsequent post-hoc comparisons, corrected for multiple comparisons using the False Discovery Rate (FDR), further delineated these effects. Pair-wise comparisons revealed a greater positivity in the difference wave (negative minus positive feedback in the feedback window) for the Social Reliable condition in frontal (β =1.3994, se = 0.202, z = 6.92, p<.0001), and central (β =0.7101, se = 0.335, z = 2.686, p<.0073) ROIs, but not in the parietal ROI (β =0.0912, se = 0.247, z = 0.369, p=0.7124) (Table 7). These results show a, wider, more frontal, distribution for the Social Reliable compared to the Symbolic Reliable condition

**Table 7.** Post-hoc comparisons for topographical distribution of feedback difference waves.

| **contrast** |  | **estimate** | **SE** | **df** | **z.ratio** | **p value** |
| --- | --- | --- | --- | --- | --- | --- |
| Social Rel -<br>Symbolic Rel | Frontal | 1.3994 | 0.202 | Inf | 6.929 | <.0001* |
| Social Rel -<br>Symbolic Rel | Central | 0.7101 | 0.335 | Inf | 2.686 | <.0073 |
| Social Rel -<br>Symbolic Rel | Parietal | 0.0912 | 0.247 | Inf | 0.369 | 0.7124 |

##### 3.2.2.4 Linear mixed-effects models

We ran linear mixed effects models, which included Condition, Exposure and Valence, and compared the mean voltage amplitudes in the time windows established by the permutation tests, at a subset of centro-parietal electrodes (C3, C4, CP1, CP2, Cz, Oz, P3, P4, PO3, PO4, Pz). Once again, we used the forward selection strategy to select the model that best fitted the feedback data. Model 0 included only the random intercepts. We then added the predictor Condition, then the predictor Valence (Positive feedback, Negative feedback), then the predictor Exposure, and finally their interaction (Table 8). The final lmer model [MV ∼ condition*valence*exposure + (1|participant), p= < 2.2e-16 ***, AIC=58733, LL=-29327] revealed significant interactions (Table 9).

**Table 8.** Model comparison for feedback EEG.

| Sampling Units |  | Number of total observations = 10263<br>Number of participants = 27, Number of items = 90, Number of electrodes: 11 |  |  |  |  |  |  |  |  |  |
| --- | --- | --- | --- | --- | --- | --- | --- | --- | --- | --- | --- |
| Model specification | Model name | Nested/<br>simpler<br>model | Fixed<br>effects<br>added | Random effects |  | Model fit |  |  |  | LRT test<br>against<br>nested |  |
| | | | | Participant<br>s | Electrodes | AIC | BIC | LL | df | d<br>f | $\chi^2$ |
| RE only | Null | --- | --- | Intercepts | Intercepts | 6371<br>0 | 63739 | -31851 | 4 | -<br>-<br>- | --- |
| FE main<br>effect | Condition | Null | Condition | “ | “ | 5990<br>6 | 59950 | -29947 | 6 | 2 | < 2.2e-16<br>*** |
| FE main<br>effects | Condition<br>+ Valence | Condition | +Valence | “ | “ | 5923<br>8 | 59289 | -29612 | 7 |  | < 2.2e-16<br>*** |
| Two-way<br>interaction | Condition<br>* Valence | Condition<br>n+<br>Valence | *Valence | “ | “ | 5922<br>0 | 59286 | -29601 | 9 | 2 | 1.619e-05<br>*** |
| Two-way<br>interaction | Condition<br>* Valence<br>+<br>Exposure | Condition<br>n*<br>Valence | +<br>Exposure | “ | “ | 5850<br>3 | 58604 | -29237 | 14 | 5 | < 2.2e-16<br>*** |
| Two-way<br>interaction | Condition<br>*<br>Valence*<br>Exposure | Condition<br>n*<br>Valence<br>+<br>Exposure | *Exposure | “ | “ | 5822<br>1 | 58503 | - 29071 | 39 | 2<br>5 | < 2.2e-16<br>*** |

**Table 9.**
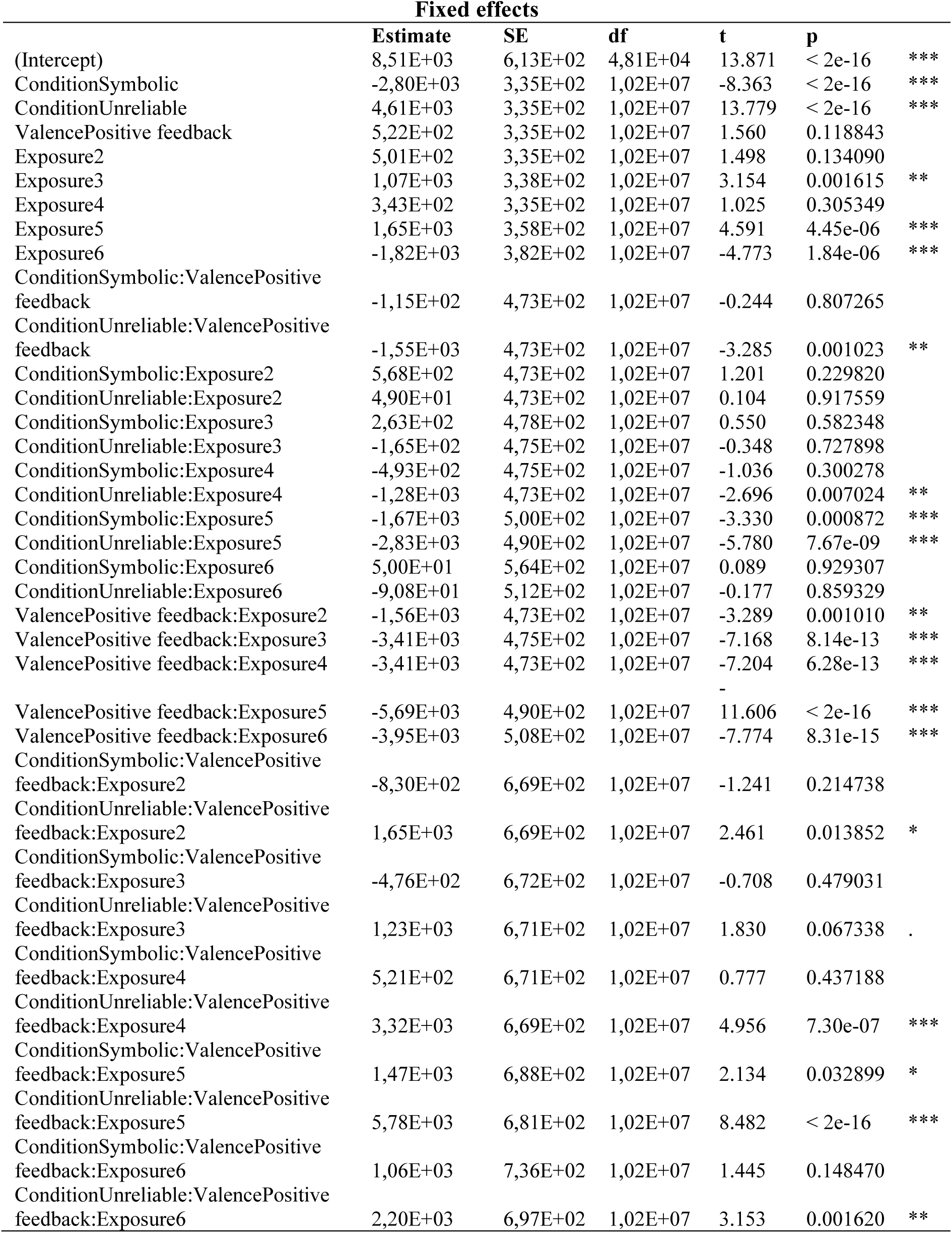
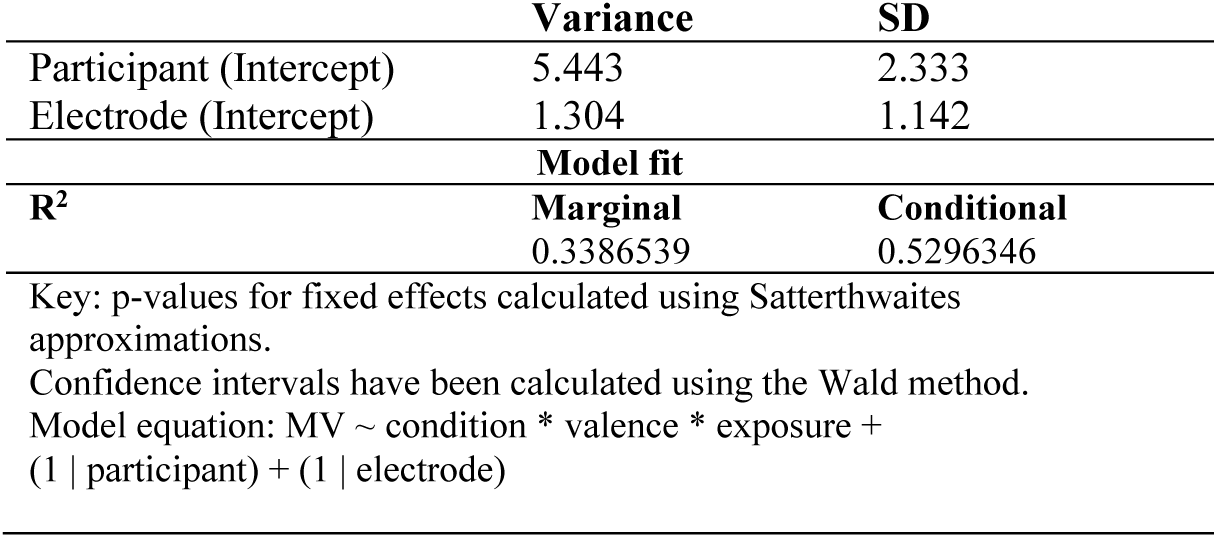
Linear fixed effects models for feedback EEG.

Subsequent post-hoc comparisons, corrected for multiple comparisons using the False Discovery Rate (FDR), further delineated these effects. Pair-wise comparisons showed that when feedback was negative, no clear pattern emerged as a function of exposure in any of the conditions. When differences between exposures were significant, these changes went in different directions, sometimes becoming more positive and others more negative (Table 10). On the other hand, positive feedback resulted in clearer pattern of decreasing positivity as the learning session progressed. This pattern was most visible in the Social Reliable condition, where a decreased positivity could be seen in five out of the five exposure changes, between exposures 1 and 2 (β =10.546, se = 0.335, z = 3.152, p<0.0019), 2 and 3 (β =12.878, se = 0.335, z = 3.850, p<.001), 3 and 4 (β =0.7226, se = 0.335, z = 2.160, p=0.0308), 4 and 5 (β =0.9790, se = 0.335, z = 2.927, p=0.0037), and 5 and 6 (β =17.251, se = 0.335, z = 5.157, p<.001).

**Table 10.** Post-hoc comparisons for feedback EEG.

Social Reliable Negative
| contrast | estimate | SE | df | z.ratio | p value |
| --- | --- | --- | --- | --- | --- |
| Exposure1-Exposure2 | -0.5012 | 0.335 | Inf | -1.498 | 0.1547 |
| Exposure2-Exposure3 | -0.5643 | 0.338 | Inf | -1.670 | 0.1294 |
| Exposure3-Exposure4 | 0.7226 | 0.338 | Inf | 2.139 | 0.0487 |
| Exposure4-Exposure5 | -13.022 | 0.358 | Inf | -3.634 | 0.0006 |
| Exposure5-Exposure6 | 34.675 | 0.402 | Inf | 8.627 | <.0001* |

Social Unreliable Negative
| contrast | estimate | SE | df | z.ratio | p value |
| --- | --- | --- | --- | --- | --- |
| Exposure1-Exposure2 | -0.5502 | 0.335 | Inf | -1.645 | 0.1154 |
| Exposure2-Exposure3 | -0.3499 | 0.335 | Inf | -1.046 | 0.3167 |
| Exposure3-Exposure4 | 18.327 | 0.335 | Inf | 5.479 | <.0001* |
| Exposure4-Exposure5 | 0.2557 | 0.335 | Inf | 0.764 | 0.4446 |
| Exposure5-Exposure6 | 0.7248 | 0.341 | Inf | 2.123 | 0.0422 |

Symbolic Negative
| contrast | estimate | SE | df | z.ratio | p value |
| --- | --- | --- | --- | --- | --- |
| Exposure1-Exposure2 | -10.693 | 0.335 | Inf | -3.197 | 0.0021* |
| Exposure2-Exposure3 | -0.2589 | 0.338 | Inf | -0.766 | 0.5543 |
| Exposure3-Exposure4 | 14.778 | 0.341 | Inf | 4.333 | 0.0001* |
| Exposure4-Exposure5 | -0.1300 | 0.352 | Inf | -0.369 | 0.7630 |
| Exposure5-Exposure6 | 17.528 | 0.428 | Inf | 4.093 | 0.0001* |

Social Reliable Positive
| contrast | estimate | SE | df | z.ratio | p value |
| --- | --- | --- | --- | --- | --- |
| Exposure1-Exposure2 | 10.546 | 0.335 | Inf | 3.152 | 0.0019* |
| Exposure2-Exposure3 | 12.878 | 0.335 | Inf | 3.850 | 0.0001* |
| Exposure3-Exposure4 | 0.7226 | 0.335 | Inf | 2.160 | 0.0308* |
| Exposure4-Exposure5 | 0.9790 | 0.335 | Inf | 2.927 | 0.0037* |
| Exposure5-Exposure6 | 17.251 | 0.335 | Inf | 5.157 | <.0001* |

Social Unreliable Positive
| contrast | estimate | SE | df | z.ratio | p value |
| --- | --- | --- | --- | --- | --- |
| Exposure1-Exposure2 | -0.6412 | 0.335 | Inf | -1.917 | 0.0691 |
| Exposure2-Exposure3 | 19.219 | 0.335 | Inf | 5.745 | <.0001* |
| Exposure3-Exposure4 | -0.2562 | 0.335 | Inf | -0.766 | 0.5121 |
| Exposure4-Exposure5 | 0.0747 | 0.335 | Inf | 0.223 | 0.8234 |
| Exposure5-Exposure6 | 25.624 | 0.335 | Inf | 7.660 | <.0001* |

Symbolic Positive
| contrast | estimate | SE | df | z.ratio | p value |
| --- | --- | --- | --- | --- | --- |
| Exposure1-Exposure2 | 13.165 | 0.335 | Inf | 3.936 | 0.0001* |
| Exposure2-Exposure3 | 12.390 | 0.335 | Inf | 3.704 | 0.0003* |
| Exposure3-Exposure4 | 0.4808 | 0.335 | Inf | 1.437 | 0.1614 |
| Exposure4-Exposure5 | 12.038 | 0.335 | Inf | 3.599 | 0.0004* |
| Exposure5-Exposure6 | 0.4150 | 0.335 | Inf | 1.241 | 0.2147 |

The Symbolic Reliable condition also showed a similar pattern for 3 out of 5 of the exposure changes, between exposures 1 and 2 (β =13.165, se = 0.335, z = 3.936, p<.001), 2 and 3 (β =12.390, se = 0.335, z = 3.704, p=.0003) and 4 and 5 (β =12.038, se = 0.335, z = 3.599, p=0004). Finally, in the Social Unreliable condition, changes were significant for 2 out of the 5 exposure changes, between exposures 2 and 3 (β =19.219, se = 0.335, z = 5.745, p<.001) and 5 and 6 (β =25.624, se = 0.335, z = 7.660, p<.001). It should be noted that in the Symbolic Reliable condition, for positive feedback, changes between the other exposures, although not significant, showed a decreasing positivity as the learning session progressed, following the pattern of the significant changes. On the other hand, in the Social Unreliable condition, the non-significant changes showed am increasing positivity, and hence the lessening pattern was not observed in this condition.

#### 3.2.3 Correlation of different waveform during feedback (positive minus negative) and behavioral performance in post-test

A Pearson correlation was computed to determine the relationship between the amplitude of difference wave (difference in amplitude between positive and negative feedback electrode Pz) during the time windows established by cluster-based permutation tests (see *3.2.2.2*) during feedback processing and average accuracy post-training. The results indicate a significant, positive relationship between behavioral accuracy and difference wave amplitude in the Social Reliable condition (r=0.51, p=0.0052), a null correlation in the Social Unreliable condition (r=0.01, p=0.959) and a non-significant, positive relationship in the Symbolic Reliable condition (r=0.31, p=0.0972). The Fisher’s Z-Transformation test showed that the Social Reliable correlation was significantly different from that of the Social Unreliable condition (Z = 1.97, *p* = .0489). Symbolic Reliable and Social Unreliable correlations were not significantly different from each other (z=1.14, p=.2560).

### 3.3 Pupillometry

#### 3.3.1 Pre-feedback

##### 3.3.1.1 Visual comparison

We compared the pupil dilation/constriction preceding feedback, per exposure, in the three conditions. Visual inspection of the pre-feedback window revealed increasingly constriction with each exposure, in the Symbolic reliable and Social Reliable conditions, and no visible differences in the Social Unreliable condition.

##### 3.3.1.2 Mixed models

We applied linear mixed effect models to the pre-feedback pupillometry data. The forward selection strategy was used to select the model that best fitted the pre-feedback data. Model 0 included only the random intercept. We then added the predictor Condition, then Exposure and finally their interaction (Table 11). The final lmer model [Pupildillation ∼ condition + exposure + (1|participant), p= 0.03715, AIC= −1483.0 LL=751.50] revealed a main effect of condition such that the Symbolic Reliable condition showed greater dilation compared to Social Reliable condition (β =1.29, se = 8.49, t = 2.370, p=0.023538). No differences emerged between the Social Reliable and Social Unreliable conditions (β =-7.89, se = 3.74, t = −0.211, p=0. 833058). (Table 12). There was also a main effect of Exposure such that exposures 5 and 6 showed greater constriction compared to exposure 1 (β =1.37, se = 5.27, t = −2.596, p=0. 009839, for exposure 5 and β =-1.07E se = 5.28 t = −2.025, p=0.043589, for exposure 6).

**Table 11.** Model comparison for pre-feedback pupillometry.

Pre-Feedback
| Sampling Units |  | Number of total observations = 372<br>Number of participants = 21 |  |  |  |  |  |  |  |  |  |
| --- | --- | --- | --- | --- | --- | --- | --- | --- | --- | --- | --- |
| Model specification | Model name | Nested/<br>simpler<br>model | Fixed<br>effects<br>added | Random effects |  | Model fit |  |  |  | LRT test<br>against nested |  |
|  |  |  |  | Participant<br>s |  | AIC | BIC | LL | df | df | p |
| RE only | Null | --- | --- | Intercepts |  | -1469.1 | -<br>1457.3 | 737.5<br>6 | 3 | --<br>- | --- |
| FE main<br>effect | Condition | Null | Condition | “ | “ | -1481.2 | -<br>1461.6 | 745.5<br>8 | 5 | 2 | 0.000326<br>3 *** |
| **FE main**<br>**effects** | **Condition+**<br>**Exposure** | **Condition** | **+Exposure** | “ | “ | **-1483.0** | **-**<br>**1443.8** | **751.5**<br>**0** | **1**<br>**0** | **5** | **0.03715**<br>**\*\*\*\*** |
| FE main<br>effects | Condition<br>*Exposure | Condition<br>+Exposure | *Exposure | “ | “ | -1472.6 | -<br>1394.2 | 751.5<br>0 | 2<br>0 | 1<br>0 | 0.4758 |

RE= Random effects; FE= Fixed effects

**Table 12.**
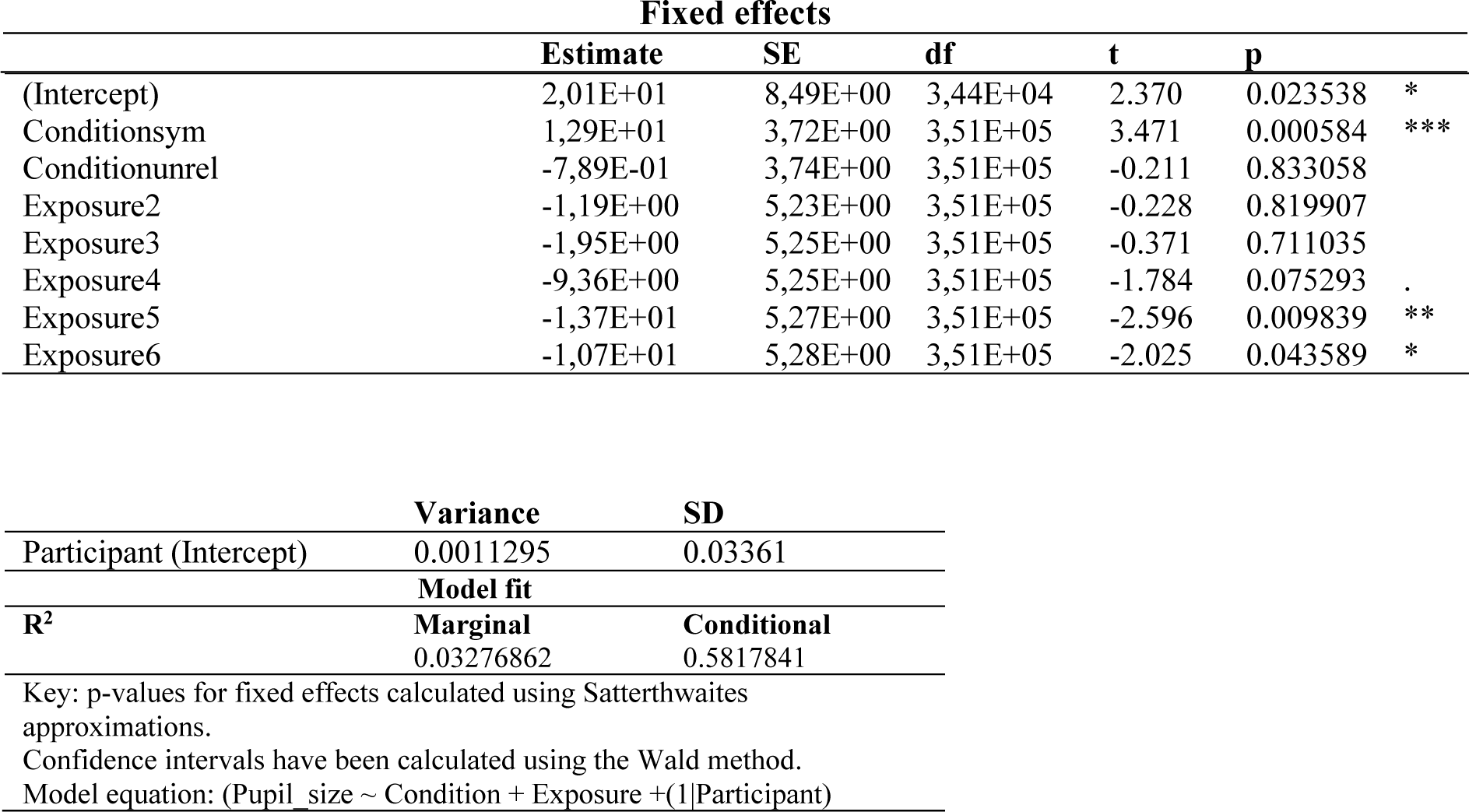
Linear fixed effects model for pre-feedback pupillometry.

#### 3.3.2 During feedback

##### 3.3.2.1 Visual inspection and Cluster-based permutation tests

As with the ERP analyses, we plotted negative and positive feedback separately for each condition. Visual inspection revealed greater dilation in the feedback window for negative compared to positive feedback in the Social Reliable and Symbolic Reliable conditions, but not in the Social Unreliable condition. Notice also the earlier effect on pupil dilatation for the symbolic compared to the social condition, due to the nature of the feedback.

Following visual inspection, we used cluster-based permutation tests to determine the time windows to be used for subsequent mixed models’ analyses. Significant clusters (shown in grey, Fig.8) emerged in the Social Reliable condition between 52 - 2000 ms (t-mass: −6286.2, p= .0011) post-stimulus, and in the Symbolic Reliable condition between 199 ms - 501 ms, (t-mass: −2338.6, p=.0309) post-stimulus. No significant clusters emerged in the Social Unreliable condition (Fig. 8).

##### 3.3.2.2 Linear mixed-effects models

We ran linear mixed effects models to compare pupil dilation using the time windows where significant positive vs negative feedback differences came out in the permutation tests, for the Social Reliable and Symbolic Reliable conditions (but not for the Social Unreliable condition as it showed no significant differences in the permutation tests). The models included Condition, Exposure and Valence. We used the forward selection strategy to select the model that best fitted the feedback data. Model 0 included only the random intercepts. We then added the predictor Condition, then the predictor Valence, then the predictor Exposure, and finally their interaction (Table 13). The final lmer model [Pupildillation∼condition+valence+exposure+(1|participant), p=3.182e-06, AIC=-15114, LL=765.70] revealed a main effect Valence, indicating Negative feedback was associated with greater pupil dilation for both the Social Reliable and Symbolic Reliable conditions. There was also a main effect of Condition, showing greater dilation for the Symbolic condition overall but no interaction with Valence. Finally, the model revealed a main effect of Exposure, showing a significant difference between the first and third, fourth, fifth and sixth exposure, in both the Social Reliable and Symbolic Reliable conditions (p<0.05) indicating that pupil constriction increased as learning progressed (Table 14, Fig.9).

**Table 13.**
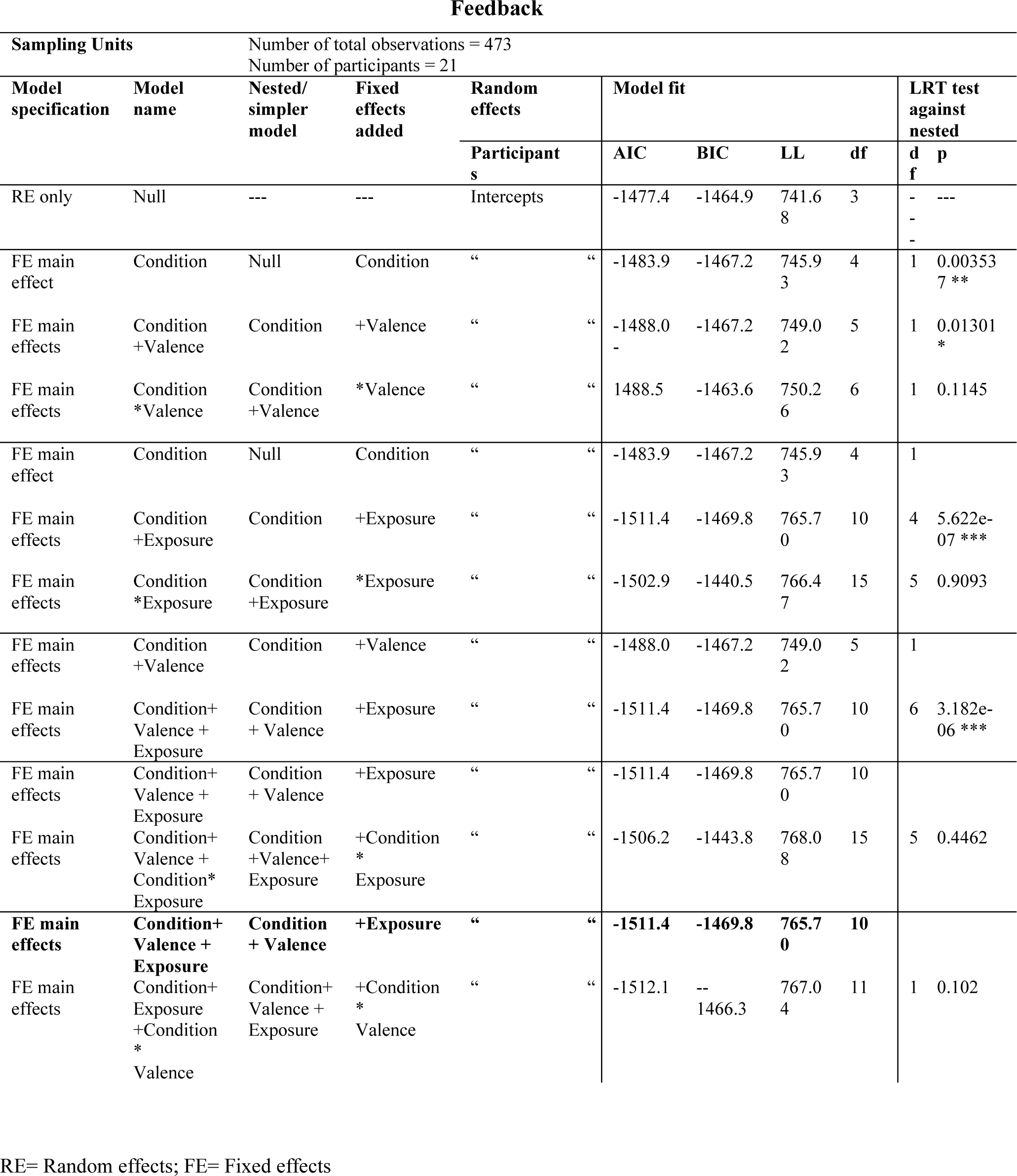
Model comparison for feedback pupillometry.

**Table 14.**
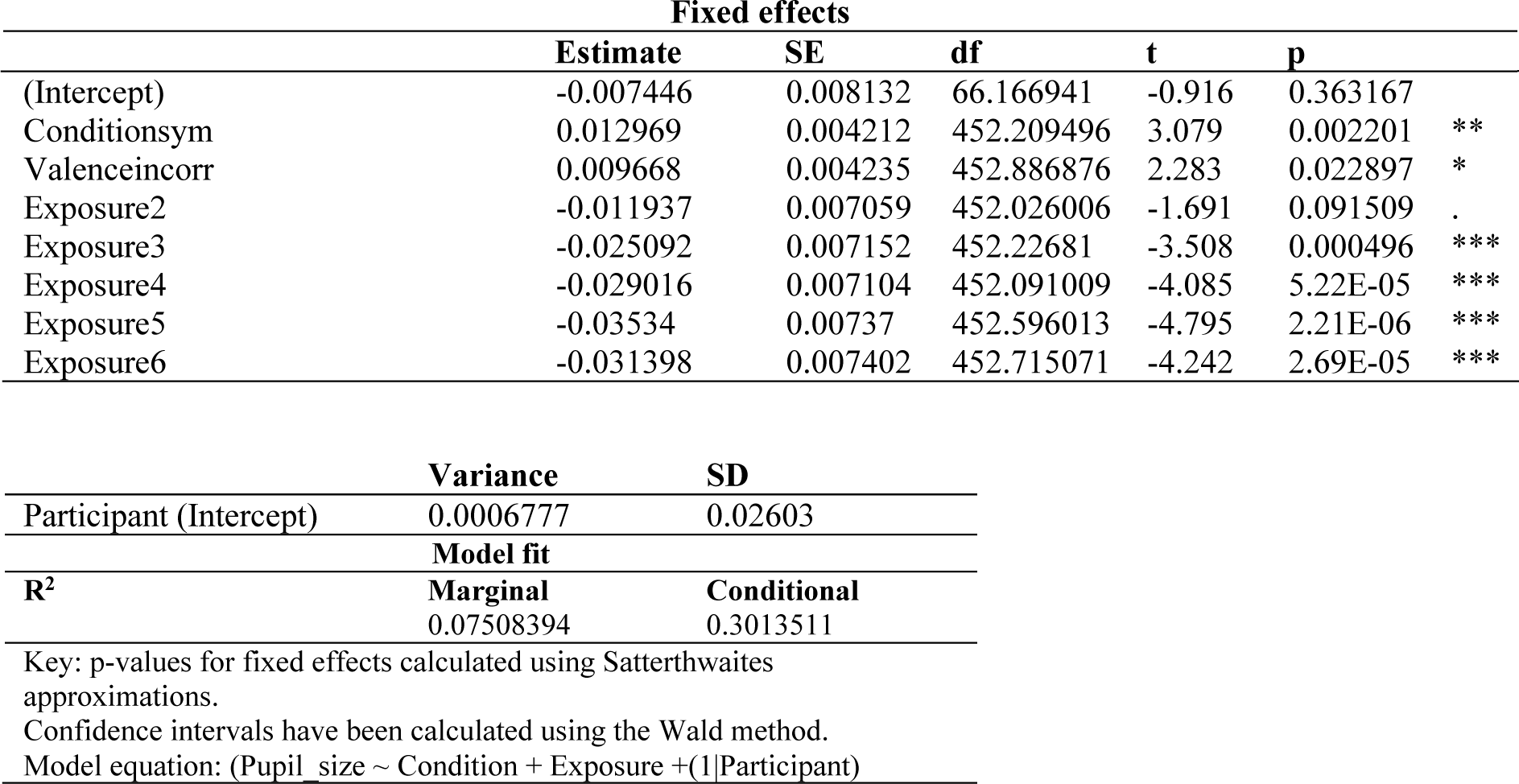
Linear fixed effects model for feedback pupillometry.

#### 3.3.3 Correlation of difference waveform during feedback (positive minus negative) and behavioral performance in post-test

A Pearson correlation was computed to determine the relationship between the size of the difference in pupil dilation between positive and negative feedback and average accuracy post-training for the Social Reliable and Symbolic Reliable conditions (during the time windows established by the cluster-based permutation tests). The results indicate a strong trend and positive relationship between behavioral accuracy and negative/positive difference in the Social Reliable condition (r=0.43, p= 0.0508), and a non-significant positive relationship in the Symbolic Reliable condition (r=0.33, p=0.1503). The Fisher’s Z-Transformation test showed that the Social Reliable and Social Unreliable correlations were not significantly different from each other (Z = 0.37, *p* = .7093).

## Discussion

In the present study, we evaluated learners’ responses to feedback with differing informational and social content during early word learning in adults. Employing a novel approach with dynamic social videos, we captured nuanced feedback expectation and processing shifts as learning progressed, via changes in ERPs and pupil dilation. Overall, our results highlight several pre- and post-feedback ERP components (i.e., stimulus-preceding negativity, SPN and a late positive complex, LPC) as key markers of feedback expectation/anticipation and processing during word learning. Crucially, using these components we showed that participants processed social feedback differently from non-social feedback. ERPs showed temporal modulations that directly reflected differences in types of feedback: an LPC effect emerged approximately at 300 ms for symbolic feedback (static still images) but was greatly delayed (app. at 1300 ms) for both social feedback conditions due to their dynamic nature (piloting data showed that it took about 1 s to interpret social feedback videos) (see Fig.4). These temporal distinctions highlight the precision with which the LPC captured feedback processing throughout word learning, offering a foundational result that guided further analyses. Additionally, this study showed effects of reliable vs non-reliable feedback in pre- and post-feedback ERP components, underscoring the importance of reliable, social feedback for word learning.

### Sensitivity to reliable vs unreliable feedback during word leaning

A comparison between how learners processed reliable versus unreliable social feedback provides valuable evidence concerning the importance of feedback informativeness during word learning. Behaviorally, learners showed improved accuracy at the end of training and in a subsequent post-test in the reliable social condition. This is in line with our hypothesis, as non-reliable (i.e., random) feedback does not allow learners to use feedback to learn; instead, they need to rely on cross-situational learning mechanisms (using statistical co-occurrences across time to infer the accuracy of the potential label-object association) (Dautriche et al., 2014; Smith & Smith, 2012; Smith et al., 2011). As outlined in the Introduction, we select information sources very carefully, prioritizing knowledgeable over inexperienced teachers (Mangardich & Sabbagh, 2018; Sabbagh & Baldwin, 2001; Scofield & Behrend, 2008: Turner et al., 2017). For instance, Mangaridch and Sabbagh (2017) showed that children disrupt semantic consolidation processes when learning from ignorant speakers as opposed to knowledgeable ones. Similarly, previous studies on adults have shown that uninformative feedback is associated with diminished SPN amplitudes compared to informative feedback (Chwilla & Brunia, 1991a; Walentowska et al., 2018). We posit that this extends to adult word learning. Unlike in the effects observed in the SPN amplitude for both reliable conditions, unreliable feedback did not show a gradual increase in the amplitude of the SPN component (see Fig.3), suggesting that unreliable feedback did not elicit increased expectations of positive feedback. Thus, these results corroborate the fact that learners are sensitive to the accuracy of information provided by feedback from potential “teachers”.

As further proof of learners’ disregard of unreliable feedback, important ERP differences emerged between reliable and unreliable feedback processing. Whereas we observed a progressive decrease in the LPC for positive feedback in the two other conditions (see Fig.5), most likely due to learners using this feedback less and less in comparison to the negative feedback, this was not the case for unreliable feedback. Also, the two reliable conditions showed significantly greater negativity in the LPC for negative vs positive feedback overall, but this was less the case in the unreliable condition. This difference likely results from learners not trusting that this feedback was indicative of their performance and not using it for context updating. Although the LPC component showed slightly more negative amplitude for negative vs positive feedback in the unreliable condition, this difference was much smaller, briefer and less widespread than that shown for the reliable conditions. Importantly, no correlation emerged between the LPC difference waveform and behavioral performance in the unreliable condition. It is therefore likely the LPC differences here are a result of an automatic recognition of negative vs positive facial expressions, and that this information was not used for context updating. Indeed, emotional information is known to elicit attention (Ohman et al., 2000), often described as *motivated attention*, or attention that is automatically allocated to emotional stimuli (Ferrari et al., 2008).

**Figure 5.**
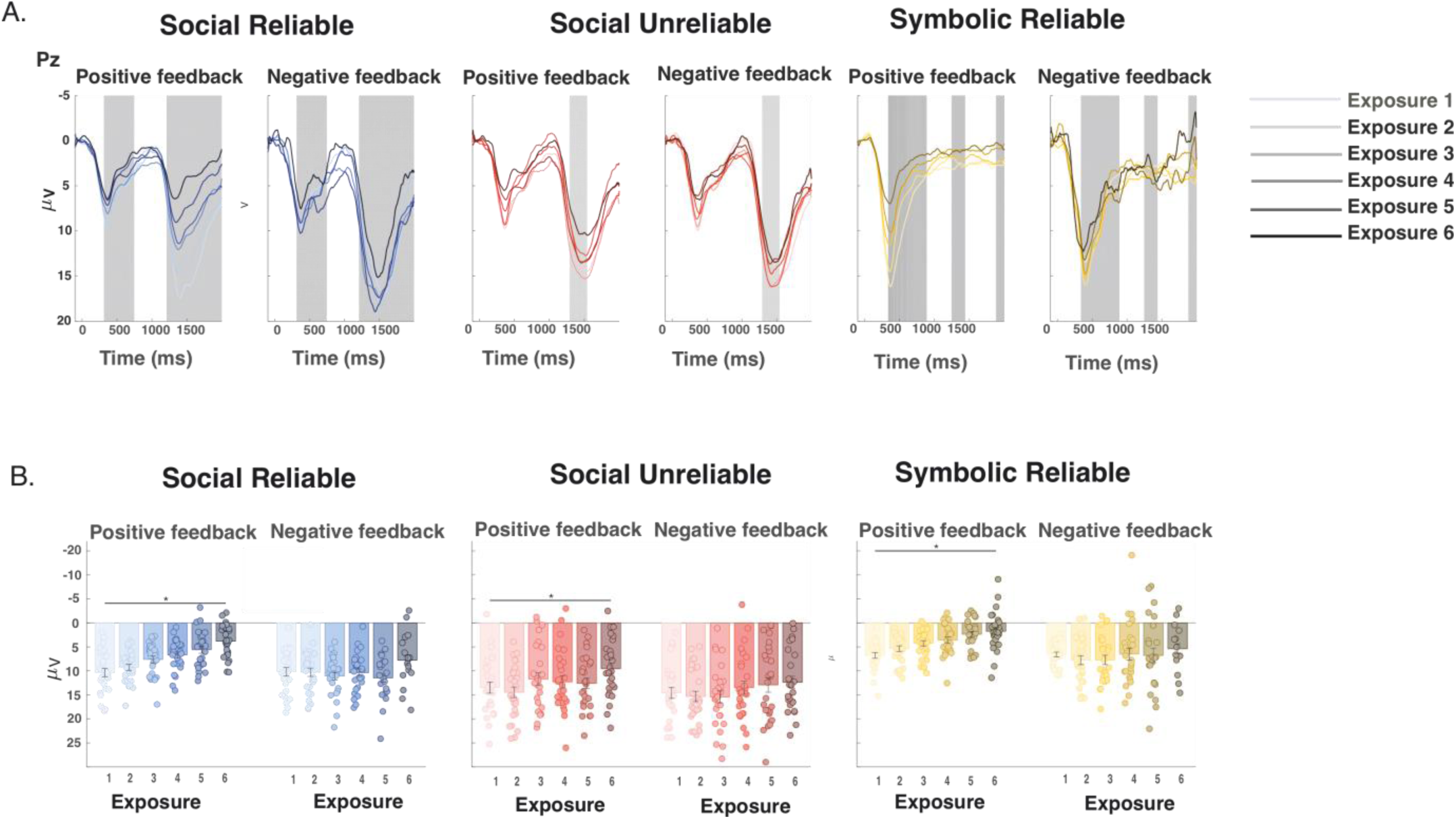
During feedback by exposure. **A.** Feedback ERPs for different learning conditions at electrode Pz and time-locked to feedback onset. For each plot, the lightest line of the given color scheme represents the 1^st^ exposure and gets progressively darker for each exposure. The grey areas show the time periods during which cluster-based permutation tests revealed significant differences between amplitudes associated with Positive and Negative feedback. **B.** Barplots represent the grand average amplitude in the analyzed time window while dots represent each participant’s mean amplitude, for the given exposure (1-6). Asterisks represent significant differences compared to the 1^st^ exposure (p<.05).

**Figure 6.**
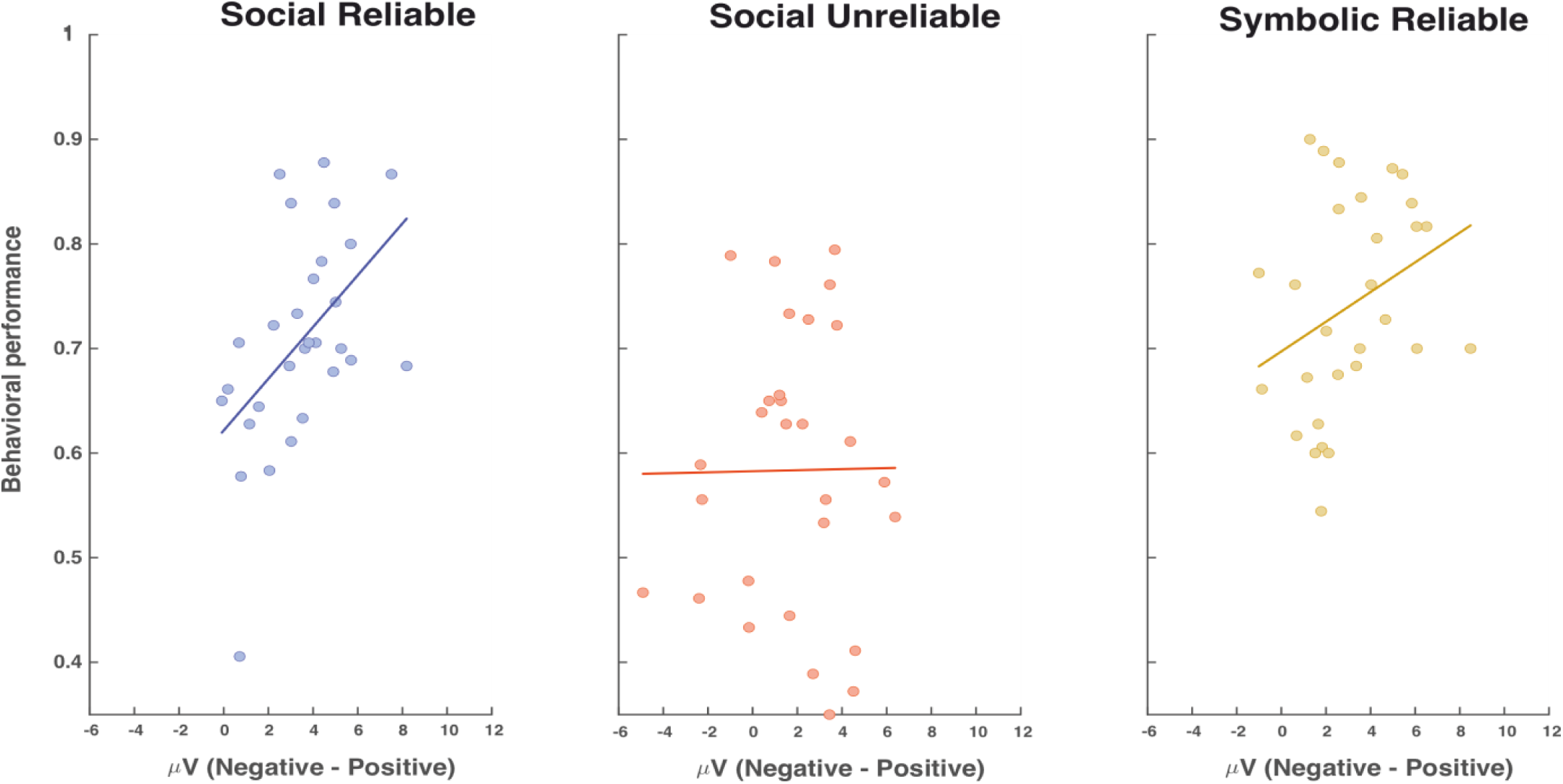
Correlation between performance post-training and negative minus positive difference wave for ERPs at electrode Pz during feedback. The x-axis represents the average amplitude of the difference waveform (negative minus positive feedback) and the y-axis represents the behavioral performance score. Each point represents a participant.

**Figure 7.**
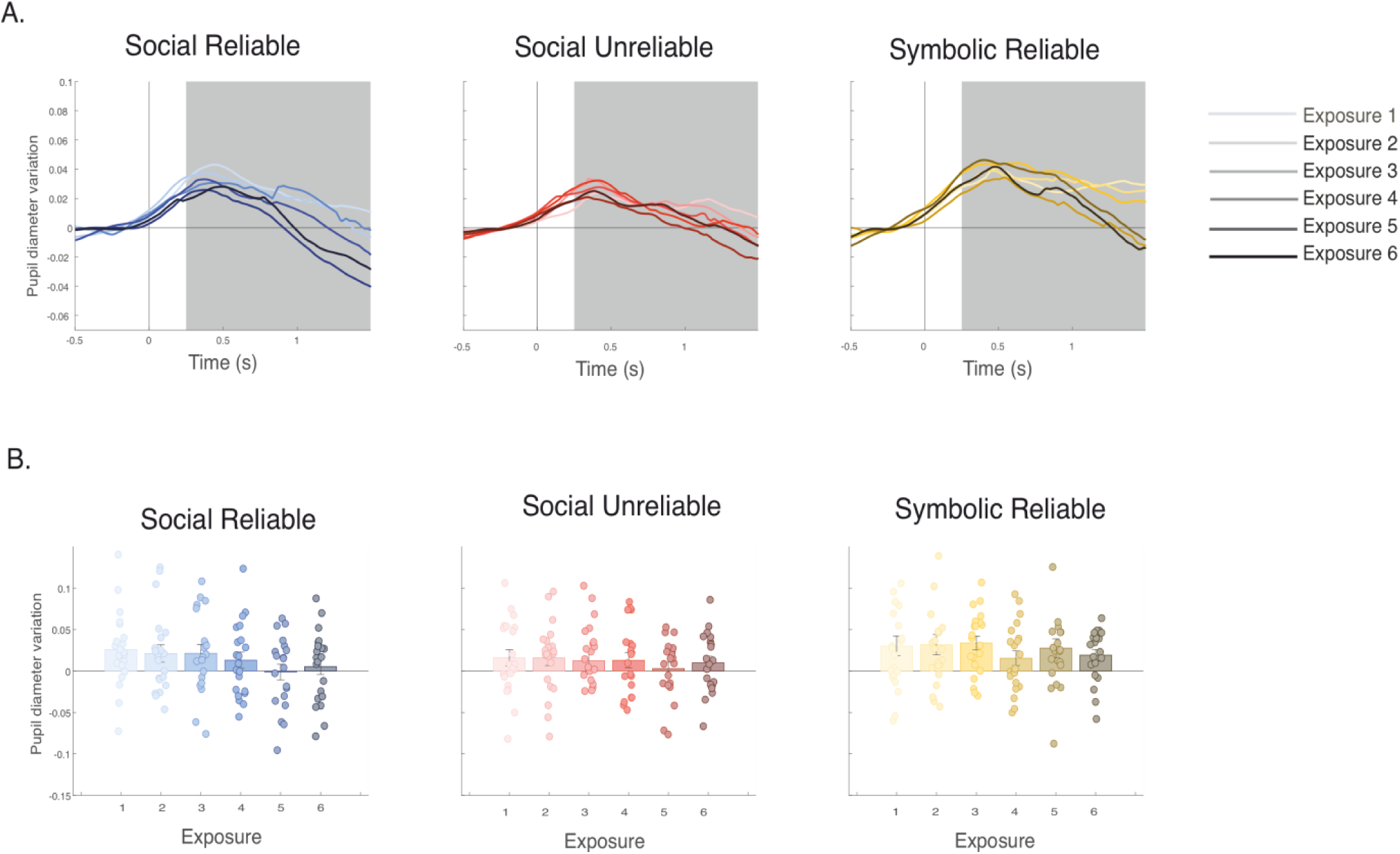
Pre-feedback by exposure. **A.** Pre-feedback pupil diameter variation (negative values represent constriction and positive values represent dilation) for different learning conditions, time-locked to the participant’s answer. Like the ERP results, for each plot, the lightest line of the given color scheme represents the 1^st^ exposure and gets progressively darker for each exposure. The grey area shows the analyzed time window (200-1000 msec). **B.** Barplots represent the grand average in the analyzed time window while dots represent each participant’s mean pupil change, for the given exposure (1-6).

Interestingly, post-training performance for words learned with unreliable feedback was above chance. It thus seems that uninformative social feedback can be overridden, and that learners can use different strategies to acquire image-word associations. Learners may have partially used implicit learning mechanisms, often employed in environments where explicit feedback is unavailable or untrusted, enabling the formation of associations through repeated exposure and statistical regularities (Morgan-Short et al., 2012). Hence, our results showing successful learning with unreliable feedback align with evidence that cross-situational statistical learning can serve as an alternative strategy in word acquisition (Angwin et al., 2022; Dautriche et al., 2021; Smith & Smith, 2012). In this sense this study also provides new evidence on how different ERP components are involved in associative and cross-situational learning (Benitez & Li, 2024; Smith et al., 2011), suggesting the implication of different cognitive processes contingent on the type of learning.

### Reliable social and nonsocial (symbolic) feedback: similarities and differences

In both the social and symbolic reliable conditions, the SPN component associated to feedback expectation increased gradually during learning (see Fig.3). Given that behavioral performance improved consistently, we propose that this reflects an increasing expectation of positive feedback. This aligns with prior research demonstrating a clear association between the SPN and learning (Moris et al., 2013; Fuentemilla, et al., 2013; Perfetti et al., 2005; Mangels et al., 2006) and the SPN and reward expectation (Fryer et al., 2021; Hackley et al., 2014; Ohgami et al., 2004). These results provide novel evidence linking the growth of the SPN during learning to positive feedback anticipation, establishing it as an important neural correlate of word learning. Concurrently, larger pupil dilation was observed for symbolic compared to social feedback conditions in the pre-feedback phase (see Fig.7). We interpret this finding as possibly reflecting the more immediate interpretability of symbolic feedback (still images), whereas social feedback (videos) required additional processing time to infer valence. Increased pupil dilation might, therefore, reflect the anticipation of more immediate feedback, though further studies are necessary to clarify how pupil responses signal feedback anticipation during learning.

During feedback processing, we showed an increase in the LPC component for negative compared to positive feedback, an effect that was observed across both reliable conditions (Donchin and Coles, 1998; Fields, 2023; Polich & Kok, 1995) (see Fig.4). This increased amplitude for negative valanced feedback is associated information-updating mechanisms and suggests a need to reevaluate the information acquired. Interestingly, the amplitude of the LPC in response to positive feedback decreased with repeated exposure (see also Luque et al., 2012) (see Fig.5). This likely reflects the diminishing utility of positive feedback as it became expected and less impactful for guiding learning. In contrast, the amplitude of the LPC for negative feedback remained steady across exposures, underscoring its ongoing relevance for information updating and learning (Bultena et al., 2017; Donchin and Coles, 1998; Fields, 2023; Polich & Kok, 1995; see Polich, 2007). Positivity in this window is also associated with more infrequent stimuli, which could explain why the LPC decreased for positive feedback, which became more and more frequent with learning. Indeed, Luque and colleagues (2012) found a very similar pattern in the same time window for positive and negative feedback during associative learning, as learning progressed.

This LPC pattern aligns with our hypotheses based on predictive learning theories, which propose that learners update associations to reduce prediction errors (Luque et al., 2012) and allocate more attention to cues signaling these errors (Pearce & Hall, 1980; Wills et al., 2007). Similarly, larger LPC effects have been observed when negative feedback is provided for high-confidence errors (Butterfield & Mangels, 2003; Metcalfe et al., 2015), i.e., when learners were highly certain about their incorrect responses. This negative increase in LPC amplitude following negative feedback likely reflects the level of surprise experienced (larger prediction error) and heightened attention to correct the previously learned association. Supporting this interpretation, Pashler et al. (2005) found that negative feedback significantly improved word learning and post-training retention.

Our findings underscore the critical role of negative feedback in updating knowledge during word learning and highlight the LPC component as a valuable neural marker for tracking this process. Moreover, a significant correlation emerged between LPC differential amplitude (negative vs. positive feedback) and subsequent learning outcomes in both reliable feedback conditions, suggesting that participants who prioritized negative over positive feedback achieved better learning outcomes. Additionally, consistent with studies linking pupil dilation to mental effort (Laeng et al., 2012), pupillometry revealed greater dilation for negative feedback (see Fig.8), further indicating that this feedback was more effectively used to update information during learning. Overall, these results align with prior findings showing that larger positivity responses during learning are associated with improved information retrieval (Azizian & Polich, 2007; Dolcos & Cabeza, 2002; Fabiani et al., 1986; Karis et al., 1984; Lemhöfer et la., 2025; Turk et al., 2018). For the first time in word learning, they demonstrate that LPC amplitude within the feedback window can serve as an indicator of context updating (Fields, 2023).

**Figure 8.**
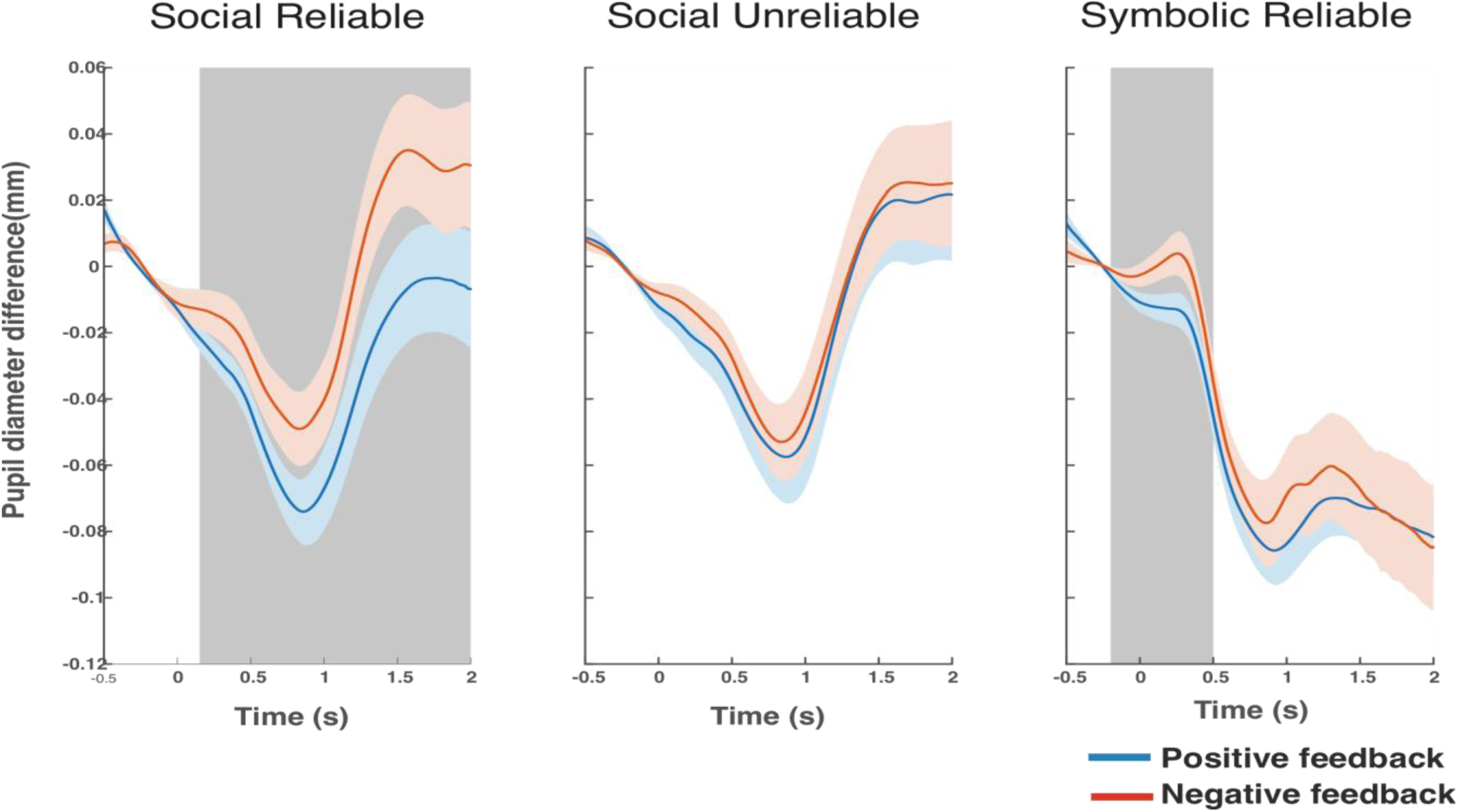
Post-feedback pupil diameter variation across learning conditions, time-locked to the feedback onset. The blue line corresponds to positive feedback and the red line to negative feedback. The grey areas show the time periods during which cluster-based permutation tests revealed significant differences between pupil variation associated with positive and negative feedback (p<.05).

**Figure 9.**
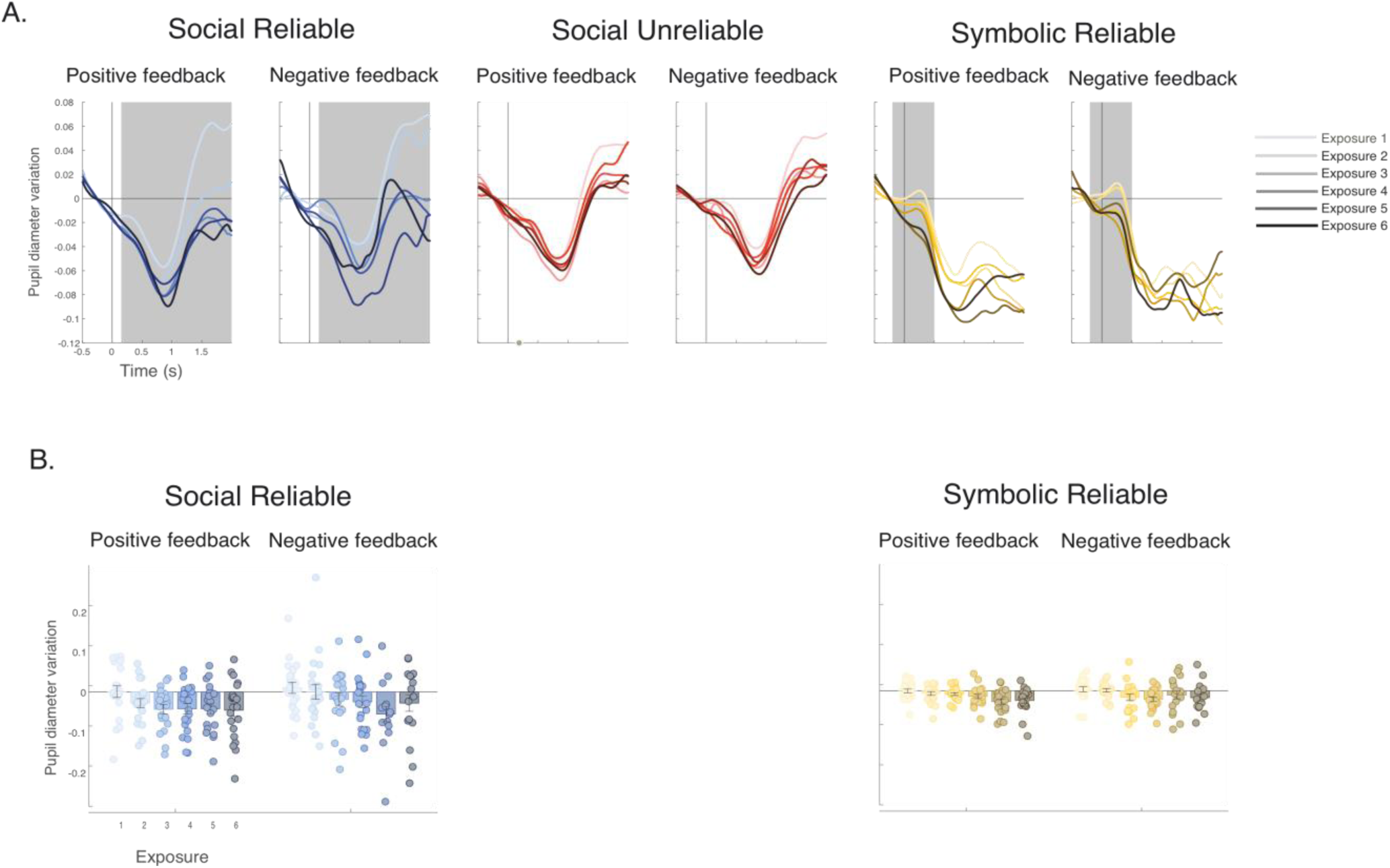
During feedback by exposure. A. Feedback pupil diameter variation (negative values represent constriction and positive values represent dilation) for different learning conditions, time-locked to the participant’s answer. Like the ERP results, for each plot, the lightest line of the given color scheme represents the 1^st^ exposure and gets progressively darker for each exposure. The grey area corresponds to the cluster-based permutation analyses (differences between pupil variation associated with positive and negative feedback). **B.** Barplots represent the grand average in the analyzed time window while dots represent each participant’s mean pupil change, for the given exposure (1-6). There is no barplot for the Social Unreliable condition as cluster-based permutation analyses did not reveal any differences between positive and negative feedback.

Regarding the differences between the reliable conditions (social and symbolic), we observed robust differences in the scalp distribution. Social feedback showed more widespread frontal central activity when compared to the symbolic condition (see Fig.4). Understanding social cues involves interpreting others’ mental states or intentions, frequently termed mentalizing or theory of mind (Fehlbaum et al., 2021; Frith & Frith, 2006). This cognitive ability is associated with the activation of a broad network, often referred to as the default mode network (DMN), including the medial prefrontal, temporoparietal junction, posteromedial cortex (Klieman & Adolphs, 2018). More specifically, Tan and colleagues (2022) recently pointed to the dorsomedial prefrontal cortex as being specialized for metalizing. Indeed, evolutionarily, the development of social cognition has been linked to the expansion of frontoparietal networks (Allen et al., 2023; Tan et al., 2022). In this context, the more widespread scalp distribution observed for our reliable social feedback, especially in more frontal regions, might suggest the engagement of mentalizing processes for interpreting social feedback. This aligns with two studies by Schindler and colleagues (2016, 2019) showing the involvement of mentalizing and “socially motivated” attention during supposed human-generated feedback processing. Like theirs, our results highlight the importance of sender identity (i.e., social vs non-social) in engaging social regions during feedback processing.

Correlations further emphasized the importance of social feedback. The strongest correlation between LPC differential amplitude (negative vs. positive feedback) and behavioral performance was observed in the reliable social condition (see Fig.6). Although this correlation was stronger in magnitude compared to the same correlation in the reliable symbolic condition, no significant differences were found between the two, limiting the interpretability of this result. Nevertheless, this observation might suggest a potential social advantage, aligning with findings by Schindler and colleagues (2022), who demonstrated that memorizing adjectives in a social condition was associated with greater LPC amplitude during encoding and improved subsequent recognition compared to control conditions. Additionally, correlation analyses revealed a positive relationship between pupil dilation and post-training accuracy (see Fig.10), but only in the reliable social condition. This finding is consistent with previous studies showing that pupil dilation during encoding predicts subsequent recognition accuracy (Papesh et al., 2012) and retrieval (Kucewicz et al., 2018; Naber et al., 2013; Pajkossy & Racsmány, 2019). Overall, the correlations observed between both LPC amplitude and pupil dilation with behavioral accuracy in the reliable social condition suggest a stronger relationship between encoding processes and word retention in social learning. This effect may reflect an optimized use of social information, consistent with prior evidence highlighting the prioritization of social cues (Wagner et al., 2016; Chevallier et al., 2012; Williams et al., 2019).

**Figure 10.**
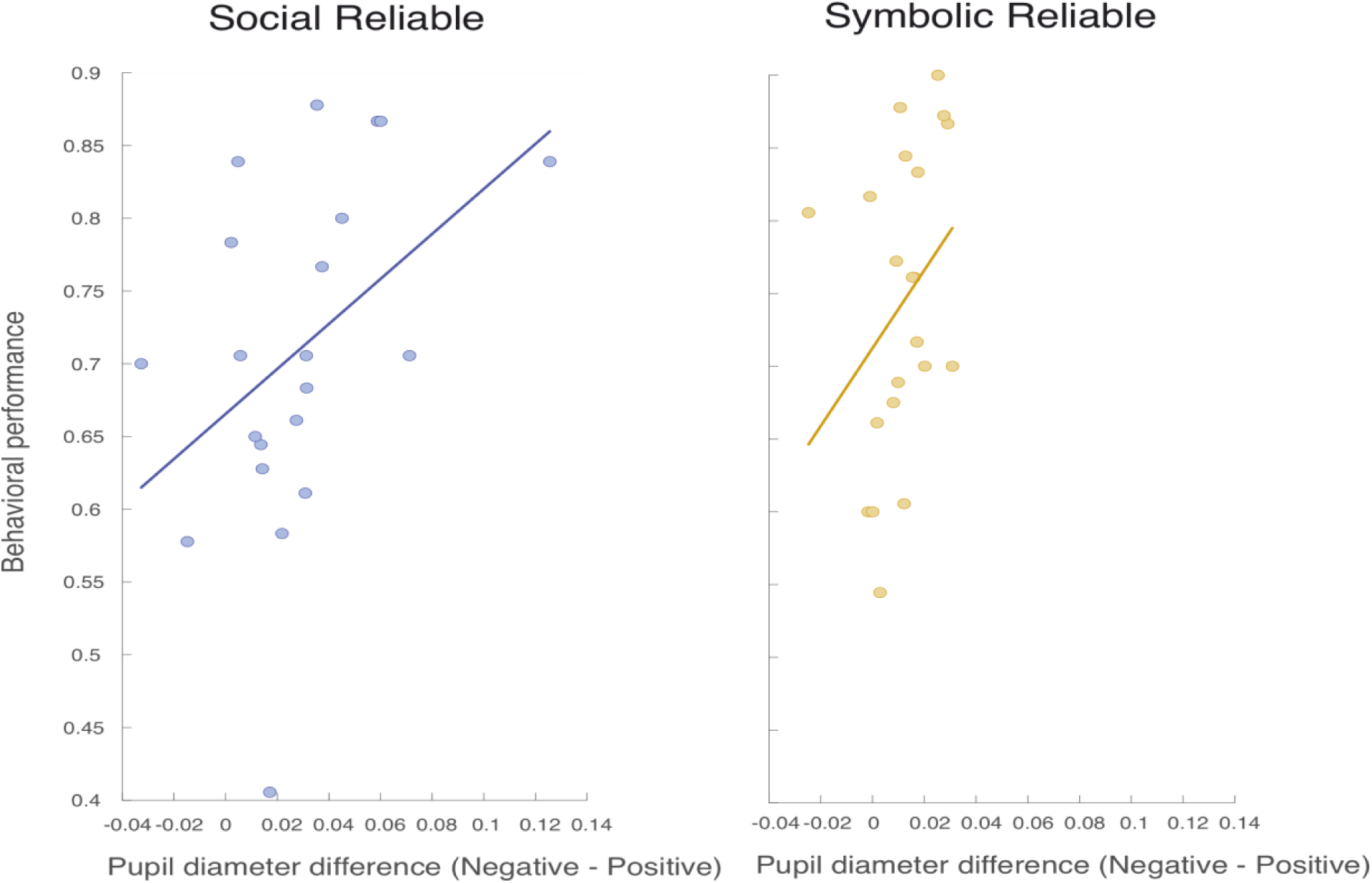
Correlation between performance post-training and pupillometry negative minus positive difference during feedback. The x-axis represents the average pupil variation of the difference waveform (negative minus positive feedback) and the y-axis represents the behavioral performance score. Each point represents a participant.

### Reliable social feedback in the context of word learning

Studies investigating how social feedback affects performance and learning have provided mixed results showing either processing advantages (Colombo et al., 2014, Pfabigan & Han, 2019; Pfabigan et al., 2018, though contingent on culture; Vernetti et al., 2017: Schindler et al., 2022) disadvantages (Beston et al., 2018, Hu et al., 2015) or no differences (Salier et al., 2024). This variability is not surprising given the diverse types of social feedback (i.e., a still image of a thumbs up, pictures of faces, dynamic videos) and the populations studied (i.e., social anxiety disorder patients, young children and adults). Furthermore, most prior studies engaged participants in tasks that were not inherently social, such as incentive delay or social incentive delay tasks (Kohls et al., 2013, Sobczak et al., 2021; Sobczak & Bunzeck, 2023, Williams et al., 2020), the probabilistic reward task (Salier et al., 2024), cue-reward association (Vernetti et al., 2017) or time estimation tasks (Pfabigan et al., 2017).

In contrast, language use - and by extension, language learning - is inherently social, making it a uniquely suitable context for investigating social feedback. Our study provides evidence that social feedback during word learning is not only behaviorally as effective as symbolic feedback, but it also exhibits reliable ERP and psychophysiological differences, pointing to the differential recruitment of cognitive processing during social feedback evaluation and retention of information. Additionally, our results highlight the critical role of feedback reliability in word learning. Unreliable feedback led to inferior behavioral outcomes and showed diminished expectation of positive feedback along with limited differentiation between positive and negative feedback during learning. It forced learners to use a different type of strategy, most probably tracking regularities across time to infer correct associations. Overall, EEG and pupillometry results, which captured feedback processing both before and during feedback, as well their relationship with performance, demonstrate that reliable social feedback shows differential processing and a tendency to have a stronger overall impact on word learning compared to symbolic feedback. Ours is the first study to examine the neural correlates of word learning with reliable social feedback, significantly advancing our understanding of how social and cognitive processes interact to shape word learning outcomes.

## Data and code availability statement

The anonymized data necessary to reproduce the analyses presented here are available on the OSF platform (https://osf.io/czhve). The materials necessary to attempt to replicate the findings presented here are not publicly accessible but are available from the corresponding author upon request.

## Authors contribution

AZ, DC and ARF planned the study. PO, MM and AZ performed the EEG measurements. XC, DC, MM, PO and AZ analyzed the data and performed the statistical analyses. AZ, DC, XC and ARF contributed to the interpretation of the data. AZ and ARF wrote the manuscript

